# Host ZCCHC3 blocks HIV-1 infection and production by a dual mechanism

**DOI:** 10.1101/2023.06.14.544911

**Authors:** Binbin Yi, Yuri L Tanaka, Hidetaka Kosako, Erika P Butlertanaka, Prabuddha Sengupta, Jennifer Lippincott-Schwartz, Akatsuki Saito, Shige H. Yoshimura

**Affiliations:** Graduate School of Biostudies, Kyoto University, Yoshida-Konoe-Cho, Sakyo-ku, Kyoto, 606-8501, Japan; Department of Veterinary Medicine, Faculty of Agriculture, University of Miyazaki, 1-1 Gakuen Kibanadai-nishi, Miyazaki, Miyazaki, 889-2192, Japan; Division of Cell Signaling, Fujii Memorial Institute of Medical Sciences, Institute of Advanced Medical Sciences, Tokushima University, 3-18-15 Kuramoto-cho, Tokushima, 770-8503, Japan; Howard Hughes Medical Institute, Janelia Research Campus, 19700 Helix Drive, Ashburn, VA 20147, U.S.A; Center for Animal Disease Control, University of Miyazaki, 1-1 Gakuen Kibanadai-nishi, Miyazaki, Miyazaki, 889-2192, Japan; Graduate School of Medicine and Veterinary Medicine, University of Miyazaki, 5200 Kiyotakecho Kihara, Miyazaki, Miyazaki, 889-1692, Japan

## Abstract

Most mammalian cells prevent viral infection and proliferation by expressing various restriction factors and sensors that activate the immune system. While anti-human immunodeficiency virus type 1 (HIV-1) host restriction factors have been identified, most of them are antagonized by viral proteins. This has severely hindered their development in anti-HIV-1 therapy. Here, we describe CCHC-type zinc-finger-containing protein 3 (ZCCHC3) as a novel anti-HIV-1 factor that is not antagonized by viral proteins. ZCCHC3 suppresses production of HIV-1 and other retroviruses. We show that ZCCHC3 acts by binding to Gag nucleocapsid protein via zinc-finger motifs. This prevents interaction between the Gag nucleocapsid protein and viral genome and results in production of genome-deficient virions. ZCCHC3 also binds to the long terminal repeat on the viral genome via the middle-folded domain, sequestering the viral genome to P-bodies, which leads to decreased viral replication and production. Such a dual antiviral mechanism is distinct from that of any other known host restriction factors. Therefore, ZCCHC3 is a novel potential target in anti-HIV-1 therapy.

---

Human immunodeficiency virus type 1 (HIV-1) infection is a major health concern, affecting tens of millions of individuals worldwide. The life cycle of the human immunodeficiency virus (HIV-1) largely relies on a large number of host proteins that promote efficient progress of viral replication, packaging, and budding. As a defence strategy, host cells produce a number of restriction factors, i.e., proteins that prevent a specific step in the viral life cycle or recognize specific viral components and activate the immune system^1, 2^. Several anti-HIV-1 restriction factors have been identified and investigated to date^3^. Their antiviral mechanism involves interaction with, modification of, and/or blocking of specific viral structures. For instance, some host factors target the viral genome. An example is the APOBEC3^4–6^ family proteins that inhibit viral proliferation by introducing nucleotide mutations in the viral genome during its reverse transcription^8, 9^. Another protein, ZAP1^10, 11^, recognizes single-stranded CG dinucleotide-rich regions of HIV-1 RNA^12, 13^ and recruits a ribonuclease and RNA exosome to degrade it. As another example, the nuclear exosome-targeting (NEXT) complex recognizes the transactivation response (TAR) element on the HIV-1 genome and inhibits transcription^14^. Other host factors target viral proteins. For instance, tetherin anchors the HIV-1 particle to the host plasma membrane and prevents the release of the virion into the extracellular space^18, 19^. SAMHD1 prevents the reverse transcription of HIV-1 by depleting the intracellular dNTP pool^20, 21^. All these restriction factors stimulate various innate immune responses, such as induction of type I interferons (IFNs) and other pro-inflammatory cytokines, triggering ubiquitin-mediated proteolysis^24, 25^. In response, HIV-1 utilizes viral accessory proteins to evade inhibition by the restriction factors. For example, the HIV-1 protein Vif antagonizes the antiviral activity of APOBEC3^6^, and the HIV-1 protein Vpu antagonizes the antiviral activity of tetherin^17, 18^. Host factors that inhibit broad HIV-1 strains and other retroviruses but are not antagonized by viral proteins are potential attractive therapeutic targets. However, no such factors, especially that target multiple steps of HIV-1 replication, have been identified to date.

CCHC-type zinc-finger-containing protein 3 (ZCCHC3; [NP_149080.2]) is a host factor with a potential antiviral activity. It was initially identified by an interactome analysis on the basis of its interaction with HIV-1 Gag^26^ and was eventually shown to regulate the activity of the interferon-stimulated response gene (ISG), which reduced the virus replication by degrading viral RNA^27^. It functionally interacts with cGAS^28^, RIG-I^29^, and TLR3^30^, major cytoplasmic nucleic acid sensors that activate the innate immune signalling pathway. ZCCHC3 acts together with cGAS to recognize double-stranded (ds) DNA and with RIG-I and TLR3 to recognize dsRNA, although no direct physical interaction between ZCCHC3 and these proteins has been reported. Porcine ZCCHC3 also suppresses the replication of the pseudorabies virus by activating the IFN-β expression^30, 31^. Despite the accumulating evidence of the antiviral role of ZCCHC3, its functional involvement in the HIV-1 life cycle remains elusive. Elucidation of ZCCHC3’s potent anti-HIV-1 activity could contribute to developing novel therapeutic strategies. Accordingly, in this study, we investigated the interaction between ZCCHC3 and HIV-1 in detail.

## ZCCHC3 and retroviral infectivity

We first confirmed that ZCCHC3 is expressed in a wide variety of cell types, including epithelial cells and monocytes (Fig. 1a). To examine the effect of ZCCHC3 on HIV-1, we transfected Lenti-X 293T cells with pNL4-3 plasmid, which is most widely used for generating an infectious HIV-1NL4-3 strain, with or without a plasmid encoding human influenza hemagglutinin (HA)-tagged human ZCCHC3 (Fig. 1b). Human ZCCHC3 cDNA was used in all experiments unless indicated otherwise. We then harvested the released virions and tested their infectivity on TZM-bl cells. TZM-bl is a reporter cell line that expresses a luciferase reporter protein, and the expression level of luciferase depends on the viral Tat protein^32–37^. While we observed a clear cytopathic effect with the NL4-3-infected TZM-bl cells (Extended Data Fig. 1a, bottom right panel), the morphological changes of cells infected with the NL4-3 virus produced in the presence of ZCCHC3 were negligible (Extended Data Fig. 1a, bottom left panel). Further, the measurement of luciferase activity in TZM-bl cells demonstrated that the overexpression of ZCCHC3 reduced the infectivity of HIV-1 (NL4-3) by two orders of magnitude (Fig. 1c), supporting the initial observation. We observed a similar effect with various HIV-1 strains, HIV-2, and simian immunodeficiency virus in chimpanzees (SIVcpz), although the extent of suppression varied among the strains (Fig. 1c). Specifically, one of the transmitted/founder (TF) HIV-1 strains, TF2625 strain, showed higher sensitivity to ZCCHC3 suppression than the TF2626 strain (p=0.0002), suggesting that susceptibility to ZCCHC3 differs between viral strains. Collectively, ZCCHC3 strongly suppresses HIV-1 infectivity.

**Fig. 1.**
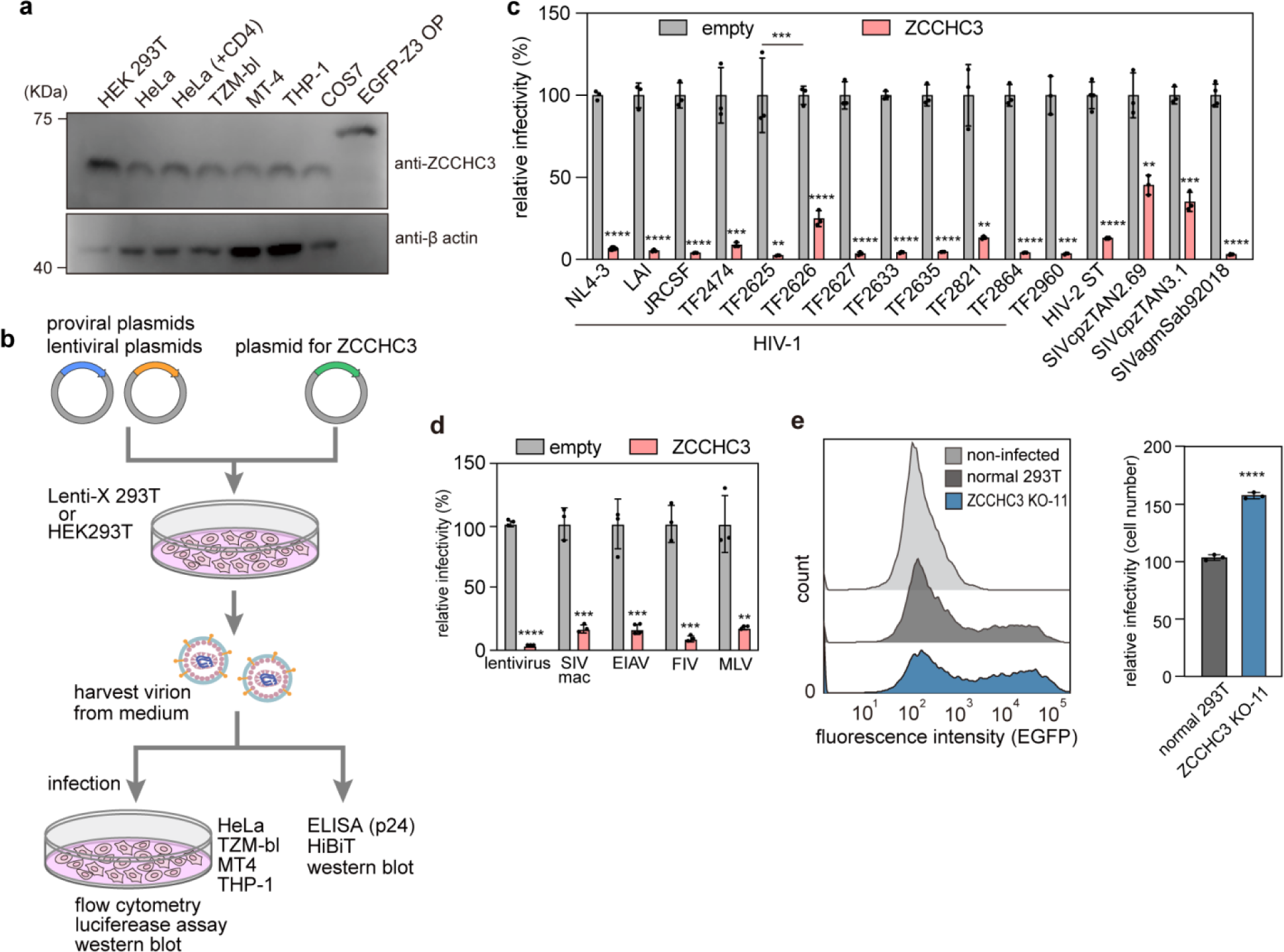
Effect of ZCCHC3 on viral infection. **a**, ZCCHC3 expression in different cell types, determined in cell lysates by western blotting with an anti-ZCCHC3 antibody. HEK293T cells expressing EGFP-tagged ZCCHC3 were the positive control. The control lane contained five times less lysate than the other lanes. The panel shown here is a representative image of three independent experiments. **b**, Experimental flow overview for panels in Figures 1 and 2. Producer cells (HEK293T or Lenti-X 293T) were transfected with viral plasmids and a ZCCHC3 expression plasmid, and the virions were collected from the culture medium and then analyzed as indicated. **c**, Effect of ZCCHC3 on the infectivity of various viral strains. Plasmids encoding individual viral strains were introduced into Lenti-X 293T cells in the presence or absence of a HA-ZCCHC3 expression plasmid. Culture supernatant was collected 2 days after transfection and used to infect TZM-bl cells. Infectivity was determined as relative light units of luciferase 2 days after infection. The mean and standard deviation values are shown (n = 3). **d**, Effect of ZCCHC3 on viral infectivity of retroviral vectors. A plasmid encoding the indicated retrovirus and luciferase reporter gene was introduced into Lenti-X 293T cells with or without a HA-ZCCHC3 expression plasmid. Culture supernatant was collected 2 days after transfection and used to infect MT-4 cells. Infectivity was determined as in (**c**). Values relative to those for cells harbouring empty vector are shown as the mean ± standard deviation. **e**, Effect of ZCCHC3 knockout on viral production. Lentiviruses produced in WT and ZCCHC3-knockout Lenti-X 293T cells were normalized to p24 and used to infect HeLa cells. The expression of a viral gene (EGFP) was assessed by flow cytometry (left). EGFP-positive cells were quantified (right); the mean and standard deviation values are shown (n = 3). Differences were examined by a two-tailed, unpaired Student’s *t*-test; \*\*\*\**p* < 0.0001, \*\*\**p* < 0.001, \*\**p* < 0.01. In panel c, difference between TF2625 and TF2626 was examined by one-way ANOVA, followed by Tukey’s test; \*\*\**p* < 0.001.

HIV-1 enters target cells via a fusion between viral Envelope (Env) and cellular membrane. To test whether the antiviral effect of ZCCHC3 is HIV-1 Env-dependent, we used a Δ*env* HIV-1 pMSMnG construct, which is pseudotyped with a VSV-G envelope, which utilizes a clathrin-mediated endocytic route to enter into cells^38^. We observed an antiviral effect of ZCCHC3 similar to that seen in the initial experiment, indicating that the antiviral effect of ZCCHC3 is HIV-1 Env-independent (Extended Data Fig. 1b). While we observed comparable antiviral inhibition by ZCCHC3 in TZM-bl and CD4^+^ T cell-derived C8166-CCR5 cells, the inhibitory effect was relatively more pronounced in THP-1 cells (Extended Data Fig. 1b), suggesting that the antiviral effect of ZCCHC3 depends on the target cell type. In addition to WT HIV-1, we tested two HIV-1 Gag capsid (CA) mutants that cannot interact with CPSF6 (N74D)^39^ or CypA (RGDA/Q112)^40^. Previous studies demonstrated that CPSF6^39^ and CypA^41^ are both involved in the early steps of HIV-1 replication, including reverse transcription, uncoating, nuclear entry, and integration targeting of proviral DNA (reviewed in ^42^). While WT and N74D viruses showed comparable sensitivity to ZCCHC3, the RGDA/Q112D virus was slightly more resistant to suppression than WT and N74D viruses (Extended Data Fig. 1c). This suggested that CA sequence may affect the susceptibility of HIV-1 to ZCCHC3.

Further, we observed that ZCCHC3 suppresses the infectivity of an HIV-1–based lentiviral vector, psPAX2-IN/HiBiT, that lacks accessory proteins, such as Vif, Vpu, Vpr, and Nef, to an extent similar to that of HIV-1 (compare Fig. 1c and Fig. 1d). This indicated that the inhibitory effect of ZCCHC3 is not antagonized by the accessory proteins. Notably, ZCCHC3 also suppressed the infection of other retroviruses, such as the simian immunodeficiency virus of macaques (SIVmac), feline immunodeficiency virus (FIV), equine infectious anaemia virus (EIAV), and murine leukaemia virus (MLV) (Fig. 1d), suggesting that ZCCHC3 inhibits the infection of a broad range of retroviruses. The suppression effect was proportional to the amount of ZCCHC3 plasmid used for viral production (Extended Data Fig. 1d). Further, *ZCCHC3* knockdown with siRNA (Extended Data Fig. 1e) and *ZCCHC3* knockout (Extended Data Fig. 1f) slightly increased lentiviral infectivity (Fig. 1e, Extended Data Fig. 1g,h). In addition, ZCCHC3 protein from different species (Extended Data Fig. 1i,j) suppressed the infectivity of JRCSF and TF2625 strains to a level similar to that of human ZCCHC3 (Extended Data Fig. 1j). The TF2626 strain was relatively resistant to ZCCHC3 from animals. The TF2625 strain, which exhibited higher sensitivity to human ZCCHC3 than TF2626 (Fig. 1c), was also more sensitive to ZCCHC3 proteins from other species tested than the TF2626 strain (Extended Data Fig. 1j). These observations suggested that the antiretroviral activity is conserved in mammals. Together, these results indicated that ZCCHC3 suppresses the infectivity of HIV-1 and other retroviruses.

## ZCCHC3 and viral production

We next assessed if ZCCHC3 decreased HIV-1 infectivity by affecting viral production. Accordingly, we quantified virions released from pNL4-3-transfected Lenti-X 293T cells into the culture medium by enzyme-linked immunosorbent assay (ELISA) of viral p24 protein. Co-expression of ZCCHC3 with HIV-1 proviral plasmids reduced the production of virions by all the viruses tested (Fig. 2a). The production was increased by 72% upon *ZCCHC3* knockout (Extended Data Fig. 2a, KO-11), and the effect was dependent on the amount of ZCCHC3 (Extended Data Fig. 2b). Western blot analysis revealed that the reduced viral production was caused by the reduction of viral protein (Gag) in the producer cells (Fig. 2b). In addition, we found that ZCCHC3 affected the processing of Gag (Fig. 2b, Extended Data Fig. 2c). Specifically, co-expression of ZCCHC3 increased the amount of p49 and p41. Together, ZCCHC3 reduced the production of virions by inducing decreased Gag expression and abnormal Gag processing.

**Fig. 2.**
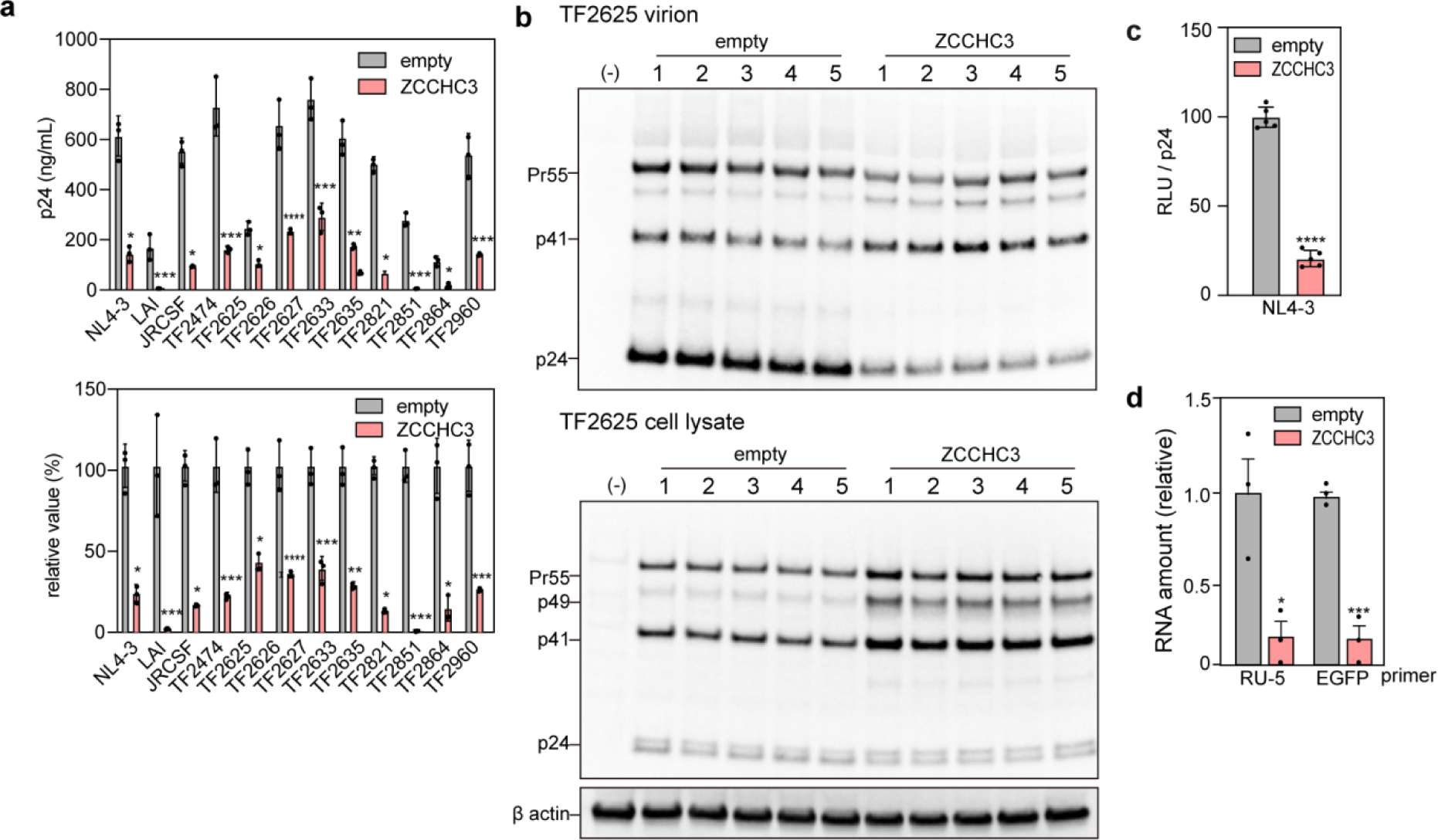
Effect of ZCCHC3 on viral production. **a**, Effect of ZCCHC3 on HIV-1 viral production. A plasmid encoding HA-tagged ZCCHC3 or empty HA-vector was introduced into Lenti-X 293T cells together with an HIV-1-encoding plasmid. Virions released into the culture medium were quantified by p24 ELISA. The absolute value (left) and the value relative to that without ZCCHC3 (right) are shown (n = 3). **b**, ZCCHC3 affects the Gag processing. Lenti-X 293T cells were co-transfected with pTF2625 plasmids in the presence or absence of HA-ZCCHC3. Equal volume of the pelleted virions and cell lysates were analyzed by western blotting using an anti-p24 antibody. The quantification of the band intensity is shown in Extended Data Fig. 2c. For cell lysate analysis, the membrane was re-probed with an anti-β-actin antibody as a loading control (n = 5). **c**, Effect of ZCCHC3 on infectivity of ZCCHC3-loaded virions. TZM-bl cells were infected with the same amount (p24-normalized) of NL4-3 or lentivirus with or without ZCCHC3 in the virion. Infectivity was analyzed by luciferase assay and is presented relative to that without ZCCHC3 as the mean and standard deviation (n = 5). **d**, Effect of ZCCHC3 co-expression on lentivirus production. HEK293T cells were transfected with lentiviral plasmids (pLV-EGFP, psPAX2, pMD2.G) with or without a HA-ZCCHC3 expression plasmid. Total RNA was purified from lentiviruses harvested from the culture medium and analyzed by RT-qPCR. The mean and standard deviation values are shown (n = 3). In panels **a**, **c, d**, differences were examined by a two-tailed, unpaired Student’s *t*-test; \*\*\*\**p* < 0.0001, \*\*\**p* < 0.001, \*\**p* < 0.001, \**p* < 0.05.

To test whether the inhibition of infection was caused by the reduced virion production, we infected TZM-bl cells with a p24-normalized virion load. Notably, NL4-3 virions produced by ZCCHC3-overexpressing Lenti-X 293T cells showed lower infectivity than those produced by non-ZCCHC3-overexpressing cells (Fig. 2c). This suggested that virions produced in the presence of ZCCHC3 are defective and have reduced infectivity. Next, to start dissecting the underlying mechanism, we analyzed the amount of viral RNA in virions. We harvested lentiviruses produced by 293T cells in the absence or presence of ZCCHC3 overexpression, and analyzed the p24-normalized virion load by reverse transcription quantitative polymerase chain reaction (RT-qPCR). Indeed, the amount of viral RNA in the virions was greatly reduced upon ZCCHC3 co-expression (Fig. 2d). Together, these results indicated that ZCCHC3 inhibits the production of infectious virus particles by both, reducing Gag expression and inhibiting the packaging of viral RNA.

## ZCCHC3 domains for GagNC–RNA binding

The described experiments suggested that ZCCHC3 directly interacts with Gag and/or viral genomic RNA. To test this, we performed a pull-down assay with different glutathione *S*-transferase (GST)-tagged Gag fragments: matrix (MAp17), capsid (CAp24), nucleocapsid (NCp7), and p6. The experiment revealed that ZCCHC3 bound to HIV-1 NCp7 and MLV GagNC (Fig. 3a). In good agreement with this, we detected ZCCHC3 in virions released into the culture medium by Lenti-X 293T cells (Fig. 3b). In fact, ZCCHC3 was present not only in TF2625 virions (Fig. 3b), but also in lentiviral virions and virus-like particles (VLPs) produced by HIV-1 Gag in the absence of other viral components (Fig. 3c). Subsequent analysis of COS7 cells expressing HA-ZCCHC3 or enhanced green fluorescent protein (EGFP)-fused ZCCHC3 and Gag-mCherry confirmed that ZCCHC3 colocalized with Gag signal both inside the producer cell and in the released VLPs (Fig. 3d). This demonstrated that a direct interaction between ZCCHC3 and GagNC is sufficient for the incorporation of ZCCHC3 into the virion.

**Fig. 3.**
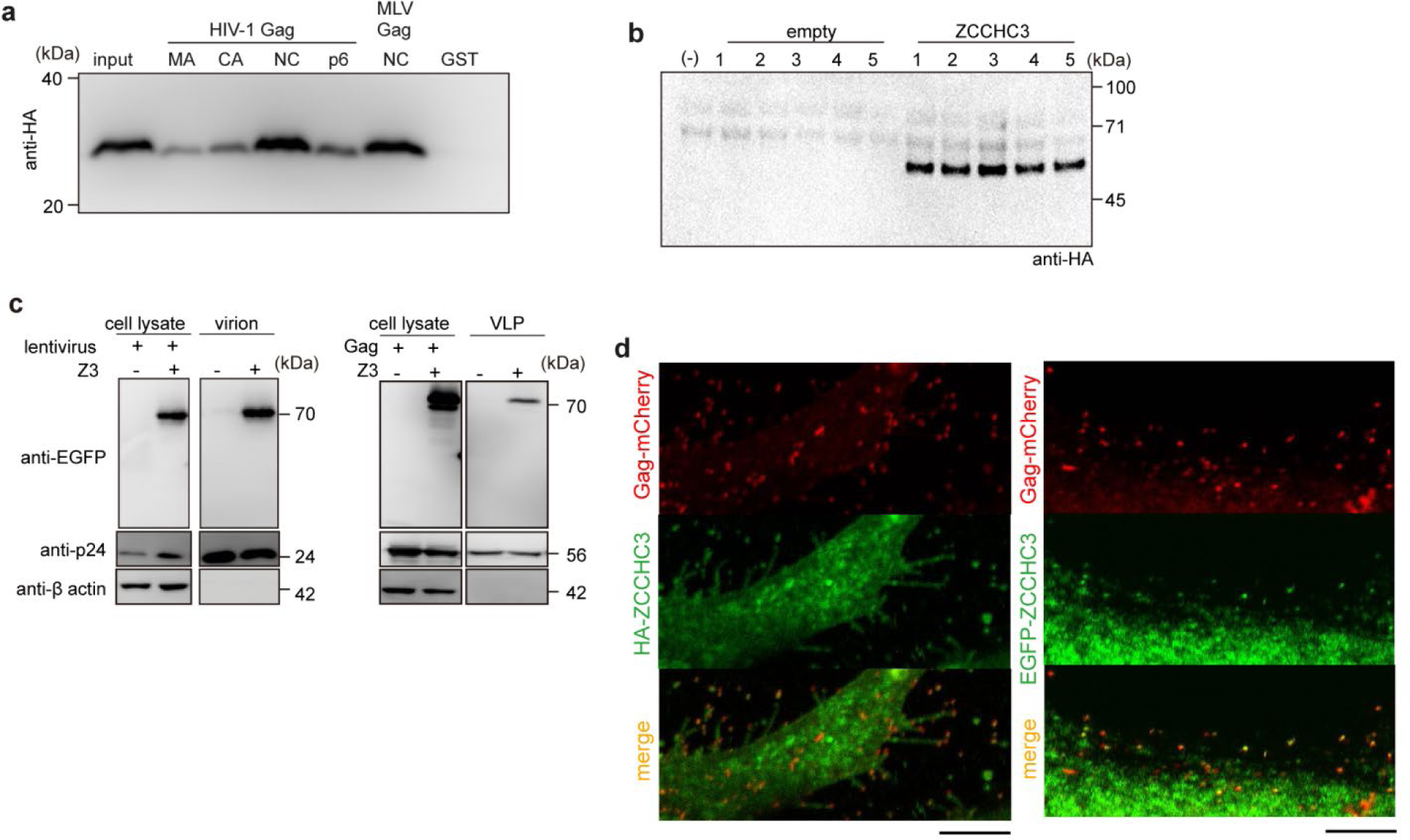
ZCCHC3 binding to GagNC. **a**, ZCCHC3 binding to NCp7 of HIV-1 Gag. GST-tagged HIV-1 Gag MAp17, CAp24, NCp7 or p6 protein, or MLV Gag NC was mixed with HEK293T cell lysate containing HA-ZCCHC3 and GSH beads. The eluted fraction was analyzed by western blotting using an anti-HA antibody. A representative image from three independent experiments is shown here. **b**, ZCCHC3 incorporated into the HIV-1 TF2625 virion. Lenti-X 293T cells were transfected with pTF2625 plasmid in the presence or absence of a HA-ZCCHC3 expression plasmid. Pelleted virions were analyzed by western blotting with a mouse anti-HA antibody (n = 5). **c**, Lentiviral plasmids (pLV-EGFP, psPAX2, and pIIIenv3-1) (left) or a plasmid encoding HIV-1 Gag (right) were introduced into HEK293T cells with or without a HA-ZCCHC3 expression plasmid. Virions were harvested by centrifugation, and analyzed by immunoblotting with anti-EGFP, anti-p24, and anti-β-actin antibodies. **d**, Presence of ZCCHC3 in HIV-1 Gag VLP. COS7 cells expressing HA-ZCCHC3 and mCherry-HIV-1 Gag were fixed, stained with an anti-HA antibody, and observed using confocal laser scanning microscopy (CLSM) (left). COS7 cells expressing EGFP-ZCCHC3 and mCherry-HIV-1 Gag were fixed and observed using CLSM (right). Scale bars, 5 µm.

We next assessed which ZCCHC3 domain is involved in the interaction with GagNC. ZCCHC3 is composed of an N-terminal intrinsically disordered region (IDR), middle fold (MF) domain, and three C-terminal tandem repeats of CCHC-type zinc-finger motifs (ZnF) (Fig. 4a). We expressed these fragments in HEK293T cells and quantified the viral production and infectivity. We observed that ZCCHC3 IDR (N) did not reduce lentiviral production nor infectivity, whereas the C fragment (MF and ZnF) suppressed both lentiviral production and infectivity (Fig. 4b,c, Extended Data Fig. 3a). We observed a similar tendency in the infectivity for other retroviruses (SIVmac, FIV, and EIAV) (Extended Data Fig. 3b). Together, these results demonstrated that the C-terminal fragment of ZCCHC3 had suppressive effect on HIV-1 and other retroviruses.

**Fig. 4.**
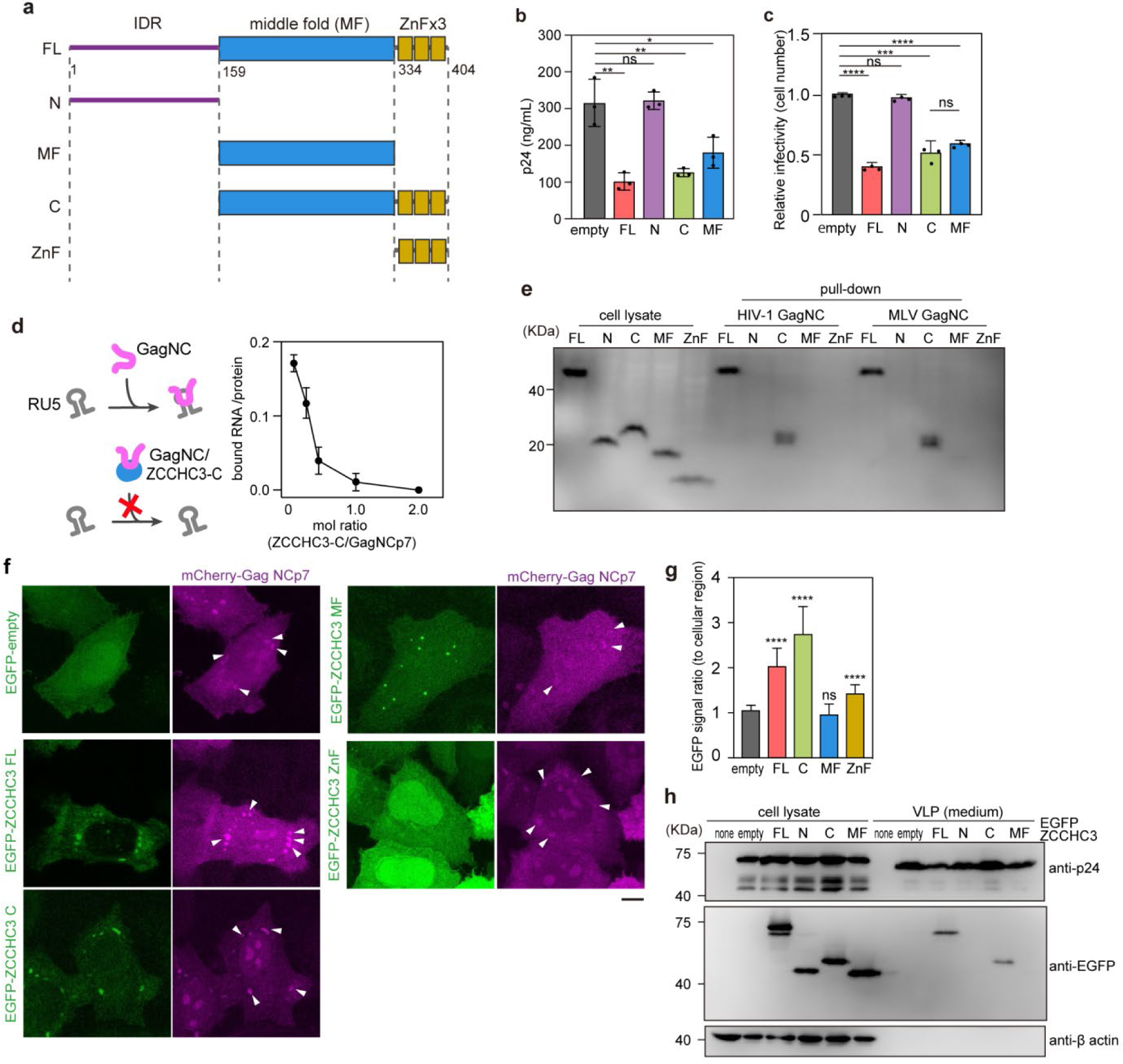
ZCCHC3 binding to GagNC via C-terminal ZnFs. **a**, The domain structure of human ZCCHC3. IDR, MF, and zinc-finger (ZnF) domains are indicated. **b,c**, Effect of C-terminal fragment of ZCCHC3 on viral production and infectivity. HEK293T cells were transfected with lentiviral vectors, and a ZCCHC3 FL, N, C, or MF expression vector or an empty vector. The resultant lentiviruses were harvested 2 days after transfection and quantified by p24 ELISA (**b**). HeLa cells were infected with a p24-normalized amount of harvested lentiviruses, and infectivity quantified based on the expression of a viral gene (EGFP) using flow cytometry (**c**). The mean and standard deviation values from three independent experiments are shown. **d**, Hypothetical mechanism of ZCCHC3 inhibition of the interaction between Gag NC and viral RNA and the effect of ZCCHC3 on the interaction between HIV-1 Gag NCp7 and LTR RNA. Purified ZCCHC3 was tested in RNA pull-down assay of Gag NCp7 and HIV-1 LTR, at the molar ratios indicated. Fluorescent probe was used to quantify RNA (ng) in the bound fraction, which is presented as a ratio to the amount of the bait protein (ng) in the bound fraction quantified by Coomassie Brilliant Blue (CBB) staining. The mean and standard deviation values from three independent experiments are shown. **e**, Binding of different ZCCHC3 domains to Gag NCp7. GST-tagged HIV-1 or MLV Gag NC was mixed with HEK293T cell lysate containing HA-tagged ZCCHC3 FL, N, C, MF, or ZnF fragments, and the eluted fraction was analyzed by western blotting with anti-HA antibody. **f,g**, Localization of HIV-1 Gag NCp7 and ZCCHC3 in HeLa cells. HeLa cells expressing mCherry-Gag NCp7 and EGFP-tagged ZCCHC3 FL, C, MF, or ZnF fragments were fixed and observed using CLSM (**f**). The arrowheads indicate the cytoplasmic GagNC condensate. Scale bar, 5 μm. The fluorescence intensity ratio of GagNC foci and cytoplasm was quantified (n = 15) (**g**). **h**, Incorporation of ZCCHC3 domains into Gag VLP. Plasmid encoding EGFP-tagged ZCCHC3 FL, N, C, or MF was introduced into HEK293T cells together with a plasmid encoding HIV-1 Gag. VLPs released into the culture medium were harvested and analyzed by immunoblotting with anti-GFP, anti-p24, and anti-β-actin antibodies. **b,c,g**, Differences were examined by a two-tailed, unpaired Student’s *t*-test; \*\*\*\**p* < 0.0001, \*\*\**p* < 0.001, \*\**p* < 0.01, ns, *p* ≥ 0.05.

Because GagNC interacts with viral genomic RNA and promotes genome packaging into the virion^33^, we speculated that ZCCHC3 inhibits the interaction between GagNC and viral RNA (Fig. 4d). Accordingly, we tested the effect of the C fragment of ZCCHC3 on RNA binding of GagNCp7 in an RNA pull-down assay. The presence of the ZCCHC3 fragment abolished the binding (Fig. 4d). This result strongly supported the notion that ZCCHC3 directly binds to GagNC. We further tested whether the amount of incorporated viral RNA affected the viral production. It is established that each viral particle contains a dimerized genomic RNA in its viral core, and the dimerized genomic RNA is critical for interaction with GagNC^41^. To this end, we produced a lentiviral vector with different amounts of transfer vector. We observed that increasing the amount of viral RNA augmented viral production (Extended Data Fig. 3c). This observation supports our result that reduction of viral RNA by ZCCHC3 in producer cells leads to decreased virion production. Together, these results demonstrated that the C-terminal fragment of ZCCHC3 inhibits the interaction between GagNC and viral RNA and reduces the viral production.

## ZCCHC3 interactions with GagNC

The C-terminal fragment of ZCCHC3 contains three ZnF motifs. ZnFs are functionally versatile structural motifs involved in various types of molecular interactions with DNA, RNA, and protein^42, 43^. We therefore investigated the role of ZnFs in the antiviral effect of ZCCHC3. A pull-down assay using purified GST-tagged GagNCp7 revealed that while the C fragment of ZCCHC3 strongly bound to GagNCp7 (Fig. 4e), the deletion of ZnFs (the MF fragment) almost completely abolished the interaction. This demonstrated that ZnFs are necessary for the interaction of ZCCHC3 and GagNCp7. We then probed ZnF-mediated interaction with Gag in cultured cells. Specifically, mCherry-fused GagNCp7 formed a liquid-like condensate in the cytoplasm (Fig. 4f, Extended Data Fig. 3d). Full-length ZCCHC3 fused with EGFP, as well as EGFP-fused C fragment of ZCCHC3, colocalized with the Gag condensate with a high partition coefficient (Fig. 4f,g). In good agreement with the pull-down assay, the deletion of ZnFs (the MF fragment) severely abrogated the colocalization, and ZnF only weakly associated with GagNCp7 (Fig. 4f,g). This demonstrated that while ZnF is necessary for the interaction of ZCCHC3 with GagNC, the MF domain is also involved in the interaction. Notably, the ZnF-mediated interaction with Gag was necessary for ZCCHC3 incorporation into the virion: the C fragment, but not the MF fragment, was incorporated into Gag VLPs (Fig. 4h) and lentiviral virions (Extended Data Fig. 3e). Together, these results indicated that ZnF is a primary site for ZCCHC3 interaction with GagNCp7, and plays a role in ZCCHC3 incorporation into the virion.

## ZCCHC3 interactions with retroviral RNA

We next examined the role of MF domain in the antiviral effect of ZCCHC3. Although, based on pull-down experiments with GST-tagged GagNCp7, the domain does not interact with Gag (Fig. 4e), it nonetheless inhibited the infectivity of HIV-1, although the effect was approximately 10% weaker than that of the C fragment (Fig. 4b,c). These observations suggested that the MF domain inhibits viral production by a mechanism distinct from the ZnF-dependent interaction with Gag.

Accordingly, we next investigated the interaction between ZCCHC3 and viral RNA. According to a structural prediction by AlphaFold2^44^, the MF domain contains a basic cleft (Fig. 5a). Further, a catRAPID prediction of RNA–protein interactions^45^ identified several potential binding sites for HIV-1 long terminal repeats (LTRs) within the MF domain (Extended Data Fig. 4a). Furthermore, our previous experiments revealed that ZCCHC3 reduces the amount of Gag in Lenti-X-293T cells (Fig. 2a,b), although the amount of viral RNA was not affected by the presence of ZCCHC3 (Extended Data Fig. 4b). Collectively, these observations suggested that ZCCHC3 functions at post-transcriptional steps of viral production.

**Fig. 5.**
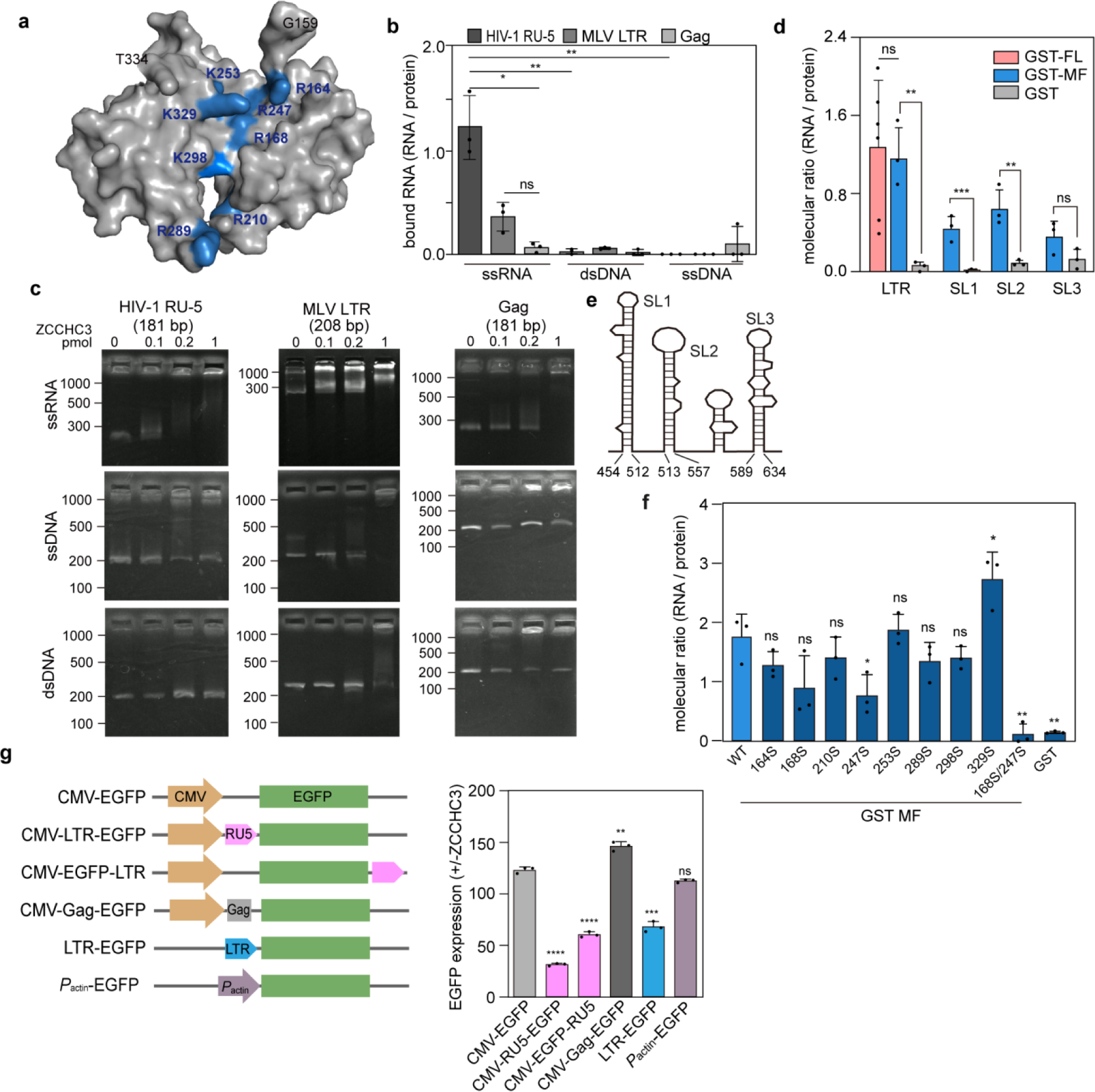
ZCCHC3 binding to retroviral RNA. **a**, 3-D structure of ZCCHC3 MF region predicted by using AlphaFold2. Basic residues in the central cleft are labelled in blue. **b**, ZCCHC3 MF domain binding to LTR RNAs of HIV-1 and MLV. GST-tagged ZCCHC3 MF or GST was mixed with ssRNA, dsDNA, or ssDNA of HIV-1 LTR (R-U5), MLV LTR, or a coding region of Gag. Nucleic acids in the bound fraction were quantified using a fluorescent probe, and the amount is presented as a molecular ratio to the bait protein. The mean and standard deviation values from three independent experiments are shown. **c**, EMSA analysis of ZCCHC3–HIV-1 genome interaction. Different amounts of purified ZCCHC3 were incubated with ssRNA, dsDNA, and ssDNA (0.1 pmol, prepared as in **b**) and analyzed. **d**, Binding of ZCCHC3 domains to LTR. RNA pull-down assay was performed with HIV-1 LTR (R-U5) and GST-tagged ZCCHC3 FL or MF as described in (**b**). The amount of bound RNA is presented as the molecular ratio to the bait protein, with the mean and standard deviation values from three independent experiments shown. **e**, Schematic illustration of HIV-1 LTR (R-U5) secondary structure. **f**, Binding of ZCCHC3 MF domain to the stem-loops of HIV-1 LTR (R-U5). RNA pull down assay was performed with the SL1, SL2, or SL3 RNA, and GST-ZCCHC3 MF (WT or R168/247S), as described in (**d**). The amount of bound RNA is presented as the molecular ratio to the bait protein, with the mean and standard deviation values from three independent experiments shown. **g**, Different EGFP-expressing constructs with the transcription of EGFP gene. HIV-1 LTR (R-U5) was inserted upstream or downstream of the *EGFP* ORF (left). A fragment of HIV-1 Gag gene (181 bp) was used as a control. Some constructs carry HIV-1 LTR (full length) or chicken β-actin promoter in place of the CMV promoter. The EGFP reporter constructs were introduced into HEK293T cells with or without a HA-ZCCHC3 expression plasmid, and EGFP fluorescence was quantified by flow cytometry. Quantification of EGFP-expressing cells as the ratio of signal between cells with (+) and without (–) HA-ZCCHC3 is shown (right). The mean and standard deviation values from three independent experiments are shown. Differences in **d** and **f** were examined by a two-tailed, unpaired Student’s *t*-test. \*\*\**p* < 0.001, \*\**p* < 0.01, \**p* < 0.05; ns, *p* ≥ 0.05. Differences in **g** were examined by a two-tailed, unpaired Student’s *t*-test, followed by Welch’s correction. \*\*\*\**p* < 0.0001, \*\*\**p* < 0.001, ** *p* < 0.005; ns, *p* ≥ 0.01.

Accordingly, we investigated the interaction between the MF domain of ZCCHC3 and viral genome. An RNA pull-down assay revealed that ZCCHC3 MF bound to R-U5 region of HIV-1 LTR and MLV LTR, but not to the coding region of HIV-1 *gag* (Fig. 5b). The protein also bound to dsDNA and ssDNA molecules carrying the same nucleotide sequences, but with a much lower affinity (Fig. 5b). We obtained similar results in electrophoretic mobility shift assay (EMSA) (Fig. 5c). Unlike the binding to GagNCp7, the MF domain was sufficient for binding to HIV-1 LTR (Fig. 5d) and to LTRs of other retroviruses (Extended Data Fig. 4c).

The LTR of HIV-1 contains three stem-loop structures (Fig. 5e). To identify the region of HIV-1 LTR that interacts with ZCCHC3 MF, we performed a RNA pull-down assay with the individual stem-loops, which revealed that ZCCHC3 MF bound to all these structures (Fig. 5d). Further, screening of ZCCHC3 MF variants with single substitutions of basic amino acid residues in the middle basic cleft (Fig. 5a) identified two residues (R168 and R247) involved in RNA binding; a single substitution (R168S and R247S) reduced the affinity for LTR RNA to approximately 50% of that of the wild type, and the double substitution (R168/247S) further reduced to approximately 15% (Fig. 5f). Together, these results demonstrated that ZCCHC3 binds to LTR RNA via its MF domain.

## ZCCHC3 and viral RNA in P-body

Previously, proteomic analysis identified ZCCHC3 in the P-body, a membrane-less organelle that sequesters various mRNAs and inhibits protein production^46^. We therefore speculated that ZCCHC3 delivers viral RNA to the P-body and thus suppresses viral protein production. Indeed, immuno-staining of HEK293T cells with ZCCHC3 and a P-body marker protein (LSM14a) revealed that endogenous ZCCHC3 was present in the P-bodies (Fig. 6a). Overexpression of ZCCHC3 increased both, the number of P-bodies in a cell and the colocalization of ZCCHC3 in P-bodies (Fig. 6a). To confirm the interaction between ZCCHC3 and the P-body, we performed proximity-dependent biotin identification of proteins (BioID analysis) using TurboID-fused ZCCHC3. We identified 610 proteins (1,228 biotinylated peptides) as spatial neighbours of ZCCHC3, including some P-body proteins (Fig. 6b). Enrichment analysis using GO terms revealed a nearly 100-fold enrichment of P-body proteins among the biotinylated proteins (Fig. 6c). Notably, lentiviral infection and the expression of LTR-carrying RNA increased the number of P-bodies and localization of ZCCHC3 in the P-bodies (Fig. 6a,d,e). Further, the MF domain, which binds to LTR RNA, was necessary and sufficient for P-body localization of ZCCHC3 (Fig. 6f). Collectively, these results suggested that ZCCHC3 recognizes LTR-carrying RNA and sequesters it to the P-body.

**Fig. 6.**
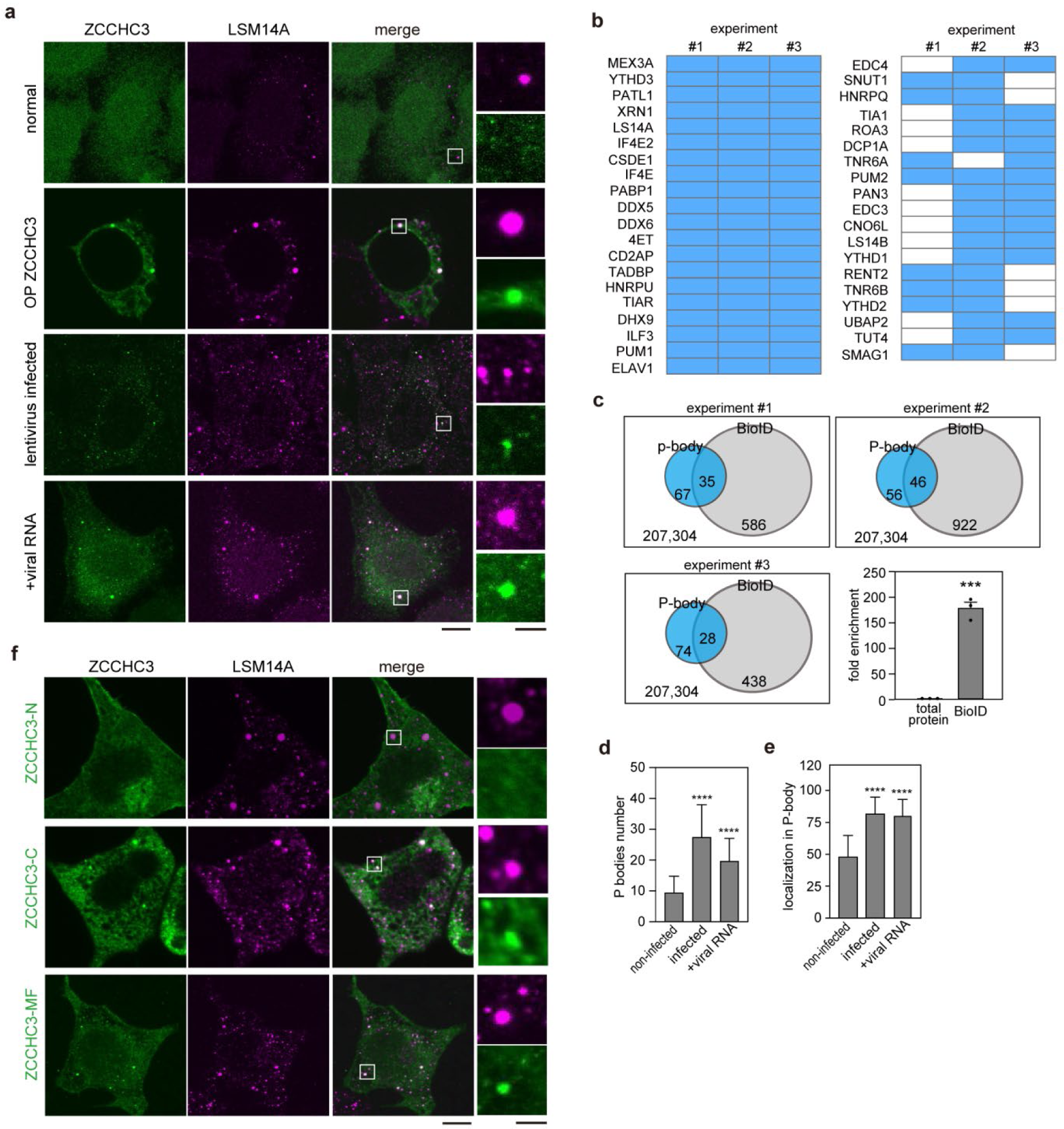
ZCCHC3 sequestration of viral RNA in P-body. **a,d,e**, ZCCHC3 localization in the P-body. 293T cells were infected with lentivirus, or transfected with a plasmid encoding EGFP-ZCCHC3 or HIV-1 LTR, and immuno-stained with anti-LSM14A and anti-ZCCHC3 antibodies. Representative images are shown (**a**). Scale bar, 5 µm. The numbers of LSM14A-positive foci per cell are shown in (**d**), with 20 cells analyzed per group. Colocalization of ZCCHC3 and LSM14A was also quantified (**e**). Differences were examined by a two-tailed, unpaired Student’s *t*-test. \*\*\*\**p* < 0.0001. **b,c**, BioID analysis of 293T cells stably expressing TurboID-fused ZCCHC3. P-body proteins were extracted from the set of identified proteins using GO terms and are shown for three independent experiments (**b**). The original data of mass spectrometry is provided in Supplementary Table 2. The total number of proteins identified in individual experiments are summarized in a Venn diagram (**c**). Fold enrichment of P-body proteins is shown as the mean and standard deviation (n = 3; **c**, right bottom panel). Differences were examined by a two-tailed, unpaired Student’s *t*-test. \*\*\**p* < 0.001. **f**, The role of ZCCHC3 MF domain in P-body localization. HEK293T cells expressing EGFP-tagged fragments (N, MF, C) of ZCCHC3 were immuno-stained with an anti-LSM14A antibody and observed using CLSM. Scale bars, 5 µm (left panels), 1µm (right panels).

## Discussion

In the current study, we demonstrated that ZCCHC3 is a novel HIV-1 restriction host factor, which acts on multiple viral components (GagNC and LTR) to suppress viral production and infectivity. We showed that ZCCHC3 binds to GagNC primarily via ZnFs and prevents it from binding to viral RNA. This results in the assembly of ZCCHC3-loaded virions, not genome-loaded virions (Extended Data Fig. 5). Further, ZCCHC3 binds to the LTR of viral genomic RNA via the basic pocket in the MF domain and sequesters the viral genomic RNA in the P-body (Extended Data Fig. 5). Notably, ZCCHC3 also suppresses a wide spectrum of retroviruses and is not antagonized by viral proteins. These unique properties make ZCCHC3 distinct from other known restriction factors.

ZCCHC3 and APOBEC3G, a well-researched HIV-1 restriction factor antagonized by viral Vif protein, are similar in that both can interact with HIV-1 genomic RNA^47^ and are incorporated in the HIV-1 virion in a GagNC-dependent manner^48^. However, the antiviral mechanism of ZCCHC3 is clearly distinct from that of APOBEC3G: ZCCHC3 sequesters the viral RNA in the P-body (Fig. 6a,b), whereas APOBEC3G increases the mutation rate at the reverse transcription step of viral life cycle^46^. Although APOBEC3G was detected in the P-body^49^, the functional significance of this observation for the antiviral effect of this protein is not known. In addition, unlike ZCCHC3 (Fig. 1c,d), APOBEC3G and other known host restriction factors are antagonized by HIV-1 accessory proteins (i.e., Vif, Vpr, Vpu, and Nef) and capsid mutations^50–52^.

We found that the ZCCHC3–viral RNA interaction is mediated by the MF domain (a.a. 159–334) but not the ZnF domain (a.a. 334–404) (Fig. 5d). The ZnF domain plays a major role in the protein’s interaction with GagNC (Fig. 4e,f,g). That was unexpected because in many host restriction factors, ZnF motifs are mainly involved in viral RNA binding^53^. Previously, ZCCHC3 ZnF was shown to bind to dsRNA and poly I:C with a K_d_ of 40 nM, although the protein fragment assayed in the previous study (a.a. 300–404) also contained a part of the MF domain^30^. Mutagenesis experiments presented herein revealed that two basic residues of ZCCHC3, R168 and R247, are involved in viral RNA binding. These two residues are predicted to be located opposite one another within the middle cleft (Fig. 5a). The predicted 3D structure of this RNA binding motif is distinct from that of any other known RNA-binding motifs. Crystal structures of ZCCHC3 MF fragment and its complex with viral RNA are needed to confirm the spatial relationship between the two identified residues, and their roles in interactions with viral RNA.

The interaction between ZCCHC3 (ZnFs) and GagNC is a novel type of protein–protein interaction. Both proteins contain CCHC-type ZnFs (3 and 2, accordingly). We observed that mCherry-fused GagNCp7 formed a liquid-like condensate in the cytoplasm when it was overexpressed in HEK293T cells (Fig. 4f, Extended Data Fig. 3d). This is in good agreement with a recent study demonstrating that GagNC undergoes liquid–liquid phase separation (LLPS) together with viral RNA, which plays a critical role in viral packaging and maturation^54^. We found that ZCCHC3 colocalized in GagNC condensate in a ZnF-dependent manner (Fig. 4f,g). This result implies that CCHC ZnFs undergo LLPS in the intracellular milieu, and that ZCCHC3 utilizes this mechanism to associate with the viral component. Indeed, recent studies demonstrated that CCHC-type ZnF, as well as other types of ZnF, undergo LLPS, both *in vivo* and *in vitro*^55, 56^. While Zn^2+^-mediated conformation of the CCHC sequence and/or electrostatic interactions among charged residues within and between individual ZnFs could be involved in the ZCCHC3–GagNC interaction, further structural and functional studies are required to elucidate the molecular details.

A limitation of this study is that most of the experiments herein used an overexpression system to test the antiviral activity of ZCCHC3. To better understand the impact of the physiological level of ZCCHC3 on HIV-1 replication, it will be interesting to use primary CD4+ T cells or macrophages for the analysis. Our data showed that the ZCCHC3-containing virus showed significantly lower infectivity in monocyte-derived THP-1 cells, suggesting that cell type may affect the antiviral activity of ZCCHC3. Furthermore, the impact of the single nucleotide polymorphism (SNP) in the *Zcchc3* gene on the anti-HIV-1 activity should be investigated. Overall, here, we demonstrated that ZCCHC3 is a novel anti-HIV-1 host factor that evades antagonization by viral proteins. Given that ZCCHC3 targets multiple steps of HIV-1 replication, it is important to test the therapeutic effect of its overexpression in cells persistently or latently infected with HIV-1. Furthermore, structural and biochemical investigations are required to design a modified ZCCHC3 with higher potency.

## Methods

### Materials

The following reagents were obtained through the NIH HIV Reagent Program, Division of AIDS, NIAID, NIH: HIV-1, Strain NL4-3 Infectious Molecular Clone (pNL4-3) (ARP-2852^57^) contributed by Dr. M. Martin, Panel of Full-Length Transmitted/Founder (T/F) HIV-1 Infectious Molecular Clones (ARP-11919^58, 59^) contributed by Dr. John C. Kappes, Human Immunodeficiency Virus Type 2 (HIV-2) ST Infectious Molecular Clone (ARP-12444) contributed by Dr. Beatrice Hahn and Dr. George Shaw, SIVcpzTAN 2.69 Infectious Molecular Clone (ARP-11497^60^) and SIVcpzTAN3.1 Infectious Molecular Clone (ARP-11498^60^) contributed by Drs. Jun Takehisa, Matthias H. Kraus and Beatrice H. Hahn, SIVagmSab92018ivTF (ARP-12140) contributed by Drs. Frank Kirchhoff, Clement Gnanadurai, and Beatrice Hahn, and SIV Packaging Construct (SIV3+, ARP-13456) and SIV LTR Luciferase mCherry Reporter Vector (ARP-13455) contributed by Dr. Tom Hope. psPAX2-IN/HiBiT and pWPI-Luc2 were kind gifts from Dr. Kenzo Tokunaga^61^. pMSMnG was a kind gift from Dr. Jun-ichi Sakuragi^62^. pLionII (1730; http://n2t.net/addgene:1730; RRID: Addgene_1730) and pCPRDEnv (1732; http://n2t.net/addgene:1732; RRID: Addgene_1732) were gifts from Dr. Garry Nolan. pEIAV-SIN6.1 CGFPW (44171; http://n2t.net/addgene:44171; RRID: Addgene_44171) and pEV53D (44168; http://n2t.net/addgene:44168; RRID: Addgene_44168) were gifts from Dr. John Olsen. lentiCRISPR v2 was a gift from Feng Zhang (Addgene plasmid# 52961; http://n2t.net/addgene:52961; RRID: Addgene_52961)^63^. pMD2.G (12259; http://n2t.net/addgene:12259; RRID: Addgene_12259) and psPAX2 (12260; http://n2t.net/addgene:12260; RRID: Addgene_12260) were gifts from Dr. Didier Trono. pLV-eGFP was a gift from Dr. Pantelis Tsoulfas (36083; http://n2t.net/addgene:36 083; RRID: Addgene_36083)^64^. pGP (# 6161) and pDON-5 Neo DNA (# 3657) were purchased from Takara Bio Inc. (Shiga, Japan). The luciferase-encoding and ZsGreen-encoding retroviral vectors were described previously^64^.

### DNA constructions

The cDNA encoding human ZCCHC3 (NM_033089.7) was amplified from cDNA pool of HeLa cells by PCR and cloned into pEGFP-C3 (Takara Bio Inc., Shiga, Japan), pCMV-HA-N2 (Takara Bio Inc.,), pET28a(+) (Takara Bio Inc.) or pGEX6P1 (Cytiva, Marlborough, MA, USA). Fragments of the ZCCHC3 N-terminus (a.a. 1 - 159), ZCCHC3 C terminus (a.a. 159-404), ZCCHC3 MF fragment (a.a. 159-334) and ZCCHC3 ZnF fragment (a.a. 334-404) were amplified by PCR using KOD-Plus-Neo (# KOD-401, TOYOBO, CO., ltd., Osaka, Japan), and subcloned into pEGFP-C3 and pCMV-HA-N2 for mammalian expression. Fragments of ZCCHC3 C terminus (a.a. 159-404) and ZCCHC3 MF fragment (a.a. 159-334) were amplified by PCR and subcloned into pET28a(+), for ZCCHC3 MF fragment also pGEX6P1 for expression in *Escherichia coli*. The cDNA encoding human cGAS (NM_138441.3) was purchased from Addgene (#108674) and subcloned into pEGFP-C3 vector.

The ZCCHC3 MF fragments carrying amino acid substitution(s) were generated by replacing the amino acids of R164, R168, R210, R247, K253, R289, K298 or K329 with S, respectively; the R268SR247S mutation was generated by replacing the amino acid of R168 and R247 with S. Single, and double amino acid mutations were introduced using PrimerSTAR® Max DNA Polymerase (# R046A, Takara Bio Inc.). The primers used for the mutation generation were listed in SI data.

The whole Gag sequence was amplified from wild-type Gag plasmid by PCR and cloned into pGEX6P1 and pmCherry-C1 (Takara Bio Inc.). Fragments of MAp17 (a.a. 1 - 132), CAp24 (a.a. 133 - 363), Gag NCp7 (a.a. 378 - 432) and p6 (a.a. 449 - 500) were amplified by PCR using KOD-Plus-Neo and subcloned into pGEX6P1 for expression in *Escherichia coli*.

The DNA fragments encoding 5’-LTR (454 - 634 nt, GenBank: MN989412.1) and Gag protein (1-181 nt) of HIV-1 were amplified by PCR using KOD-Plus-Neo and pNL4-3 as a template and cloned into pBluescript II KS(-) (212208, Agilent). The MLV 5’-LTR (1 - 207 nt, GenBank: KU324804.1) sequence was amplified from pDON-5 Neo, the EIAV 5′ LTR (1 - 114 nt, GenBank: AF247394.1) was amplified from pEIAV-SIN6.1 CGFPW, and SIV 5′ LTR (1 - 351 nt, GenBank: DQ374657.1) was amplified from SIVcpzTAN 2.69 and cloned into pBluescript II KS(-).

To generate reporter constructs, HIV-1 5’-LTR (FL) (377 - 634 nt, GenBank: MN989412.1), HIV-1 5’-LTR (R-U5) region (377 - 634 nt, GenBank: MN989412.1), a coding region of *gag* (3023 - 3203 nt, GenBank: MN989412.1), β-actin promoter on pCAGGS (386 - 661 nt) were amplified by PCR using KOD-Plus-Neo. The R-U5 fragment was cloned into NheI/AgeI or XhoI/EcoRI sites of pEGFP-C1 to generate CMV-LTR-EGFP or CMV-EGFP-LTR, respectively. The gag fragment was cloned into NheI/AgeI sites of pEGFP-C1 to generate CMV-gag-EGFP. The LTR (FL) fragment or the β-actin promoter fragment was cloned into AseI/NheI sites of pEGFP-C1 to generate LTR-EGFP or Pactin-EGFP, respectively. Information of all the primers and restriction sites for cloning is summarized in Supplementary Table 1.

### Cell culture

TZM-bl cells (ARP-8129^32–37^) were obtained through the NIH HIV Reagent Program, Division of AIDS, NIAID, NIH (contributed by Dr. John C. Kappes, Dr. Xiaoyun Wu and Tranzyme Inc.) Lenti-X 293T cells (Takara Bio Inc.), TZM-bl cells, HEK293T and COS7 cells (0539, Riken BRC Cell Bank) were cultured in Dulbecco’s Modified Eagle Medium (DMEM) high glucose (08468-16, Nacalai Tesque, Kyoto, Japan), while HeLa S3 cells (ATCC, CCL-2.2,) were cultured in DMEM with low glucose (D6046, Sigma-Aldrich, St. Louis, MO, USA), supplied with 10% (v/v) fetal bovine serum (FBS) (173012, Sigma-Aldrich). HeLa CD4 cells (NIH AIDS reagent program) were maintained in DMEM supplemented with 10% (v/v) FBS and 2 mg/mL G418 (16512-81, Nacalai Tesque, Kyoto, Japan). MT4 cells ((JCRB1216, Japanese Collection of Research Bioresources Cell Bank)), C8166-CCR5 cells, and THP-1 cells were cultured in RPMI-1640 (R8758, Sigma-Aldrich) supplemented with 10% (v/v) FBS. For passaging the adherent cells, cells were treated with 2.5 g/L-trypsin/EDTA solution (32777-44, Nacalai Tesque). All the cells were incubated in a humidified incubator at 37 °C with 5 % CO_2_.

### Plasmid transfection, virus production and collection, virus infection

Plasmid DNAs were introduced into Lenti-X 293T or HEK293T cells using either polyethylenimine hydrochloride (PEI) (49553-93-7, Polyscience, Niles, IL, USA) or TransIT^®^-293 Transfection Reagent (MIR2700, Mirus Bio LLC, Madison, WI, USA). For production of VSV-G-pseudotyped HIV-1, Lenti-X 293T cells were co-transfected with pMSMnG and pMD2.G plasmids. The supernatant was collected and filtered 48 h after transfection. For production of lentivirus, a transfer vector (pLV-EGFP or pWPI-Luc2) was introduced into HEK293T or Lenti-X 293T cells together with a packaging vector (psPAX2 or psPAX2-IN/HiBiT) and an envelope vector (pMD2.G) at a ratio of 5:3:2. The culture medium was collected 48 h after the transfection and centrifuged at 1,500 g for 10 min at 4 ℃ to remove cell debris. For other plasmid transfections, PEI was used.

Both lentiviral and retroviral vectors were rescued as described previously in the presence or absence of pCMV-HA-ZCCHC3 plasmids^65^. To rescue an SIVmac-based lentiviral vector, Lenti-X 293T cells were co-transfected with the pSIV3+ plasmid, SIV LTR Luciferase mCherry Reporter Vector, and pMD2.G plasmid. To rescue an FIV-based lentiviral vector, Lenti-X 293T cells were co-transfected with the pCPRDEnv, pLionII-luc2, and pMD2.G plasmids. To rescue an EIAV-based lentiviral vector, Lenti-X 293T cells were co-transfected with pEV53D, EIAV-SIN6.1-luc2, and pMD2.G plasmids. To rescue an MLV-based retroviral vector, Lenti-X 293T cells were co-transfected with pGP, pDON-5 Neo-luc2, and pMD2.G plasmids. The supernatant was collected and filtered 48 h after transfection.

Expression and processing of Gag proteins in Lenti-X 293T cells were evaluated with western blot. Lenti-X 293T cells transfected with either TF2625, pNL4-3 or pMSMnG were washed and lysed in 1× NuPAGE LDS sample buffer (NP0007, Thermo Fisher Scientific) containing 2% (v/v) β-mercaptoethanol and incubated at 70 °C for 10 m. For protein detection, the following antibodies were used: mouse anti-HIV-1 p24 monoclonal antibody (1:2,000, clone 183-H12-5C, ARP-3537, obtained from the HIV Reagent Program, NIH)^66^, horseradish peroxidase (HRP)-conjugated goat anti-mouse IgG antibody (1:20,000, 074-1806, KPL) and HRP-conjugated horse anti-mouse IgG antibody (1:2,000, A3854-200UL, Sigma-Aldrich). Chemiluminescence was detected using Western BLoT Ultra Sensitive HRP Substrate (T7104A, Takara) according to the manufacturer’s instruction. Bands were visualized using an iBright FL1500 Imaging System (Thermo Fisher Scientific), and the band intensity was quantified using iBright Analysis Software v5.2 (Thermo Fisher Scientific). Band intensities are presented as values relative to those without ZCCHC3.

### Purification of recombinant protein

Plasmids encoding hexahistidine (Hisx6)- and glutathione S-transferase (GST)-tagged proteins were introduced into *Escherichia coli* strain BL21(DE3) CondonPlus RIL (230245, Agilent Technologies, Inc.), and the expression of the recombinant protein was induced by 0.5 mM isopropylthio-β-D-galactoside (IPTG) in Luria Bertani broth at 18 ℃ for 6 h. The Hisx6-fused protein was purified from the cell lysate by Ni-NTA column (141-09764, Fujifilm, Tokyo, Japan) and dialyzed by 200 mM NaCl, 50 mM HEPES, 1 mM 2-mercaproethanol for 6 h at 4 ℃. The GST-fused proteins were purified by Glutathione Sepharose 4B-column (17127901, Cytiva) and dialyzed by 200 mM NaCl, 50 mM HEPES, 7:100000 (V/V) β-mercaptoethanol (99%) for 6 h at 4 ℃. The proteins were concentrated by Amicon Ultra (M.W. 3,000 Da) (Sigma-Aldrich) and stored at −80 ℃.

### Protein Pull-down assay

HEK293T cells were cultured in DMEM to 80% confluency, harvested by centrifugation at 500 g for 3 min, and re-suspended with PBS (pH 7.4) containing 1% (v/v) protease inhibitor cocktail (25955-11, Nacalai Tesque, Kyoto, Japan) 0.25% (w/v) Triton X-100 and incubated on ice for 10 min. Insoluble fraction was removed by centrifugation (1,500 g, 5 min), and the supernatant was collected. For pull-down assay using GST-fusion proteins, the purified GST-tagged Gag fragments (∼5 μg) were mixed with the lysate of HEK293T cells expressing HA- or EGFP-tagged proteins (ZCCHC3 fragments) and glutathione Sepharose™ 4B in Pull-down buffer (PBS (pH 7.4), 1 mM DTT), and incubated for 30 min at 25 ℃ with gentle rotation. The beads were washed, and the bound proteins were eluted with 200 mM glutathione in Pull-down buffer, mixed with SDS-PAGE sample buffer (100 mM Tris-HCl (pH 6.8), 4% (w/v) SDS, 20% (v/v) glycerol, 0.15 mg/mL bromophenol blue), and heated for 5 min. The proteins were analyzed by SDS-PAGE using 12% (w/v) acrylamide gel and subjected to western blot. HA- and EGFP-tagged proteins were incubated with anti-HA antibody (1: 2,000; 2999, Cell Signaling Technology, Danvers, MA, USA) or anti-EGFP antibody (1: 2,000; 598, Medical & Biological Laboratories Co., Ltd., Nagano, Japan), followed by the incubation with secondary antibody (HRP-conjugated goat anti-mouse antibody (1: 10,000; NA931V, Cytiva) or HRP-conjugated goat anti-rabbit antibody (1: 10,000; A-11036, Thermo Fisher Scientific, Waltham, MA, USA). The immunoreactive bands were visualized using Chemi-Lumi One Super Kit (Nacalai Tesque) under LAS-3000 Imager (Fujifilm). The gel was also stained with Coomassie Brilliant Blue (CBB) to check the input amount and the unbound fraction.

### RNA and DNA pull-down

GST-tagged proteins were immobilized to glutathione sepharose beads (Cytiva) and incubated with synthesized RNA for 10 min in RNA Pull-down buffer (20 mM Tris-HCl (pH 7.4), 30 mM NaCl, 0.1 mM MgCl_2_, 1 mM DTT, 1:100 RNase inhibitor (SIN-201, TOYOBO, CO., ltd., Osaka, Japan) in RNase-free water). The protein-RNA complex was eluted with 200 mM glutathione in RNA Pull-down buffer. The RNA amount in the eluted fraction was quantified with Quanti Fluor RNA dye (E286A, Promega, Madison, WI, USA) by following the manufacturer’s instruction. The eluted fraction was also subjected to SDS-PAGE, followed by CBB staining. For DNA pull-down assay, the DNA fragments were amplified by PCR using a plasmid carrying HIV-1 5′ LTR (454 - 634 nt, GenBank: MN989412.1), MLV 5′ LTR (1 - 207 nt, GenBank: KU324804.1) or Gag (935 - 1115 nt, GenBank: MN989412.1) as a template and the primers described in Supplementary Table 1. Single stranded DNAs were generated by heating the dsDNA at 95 ℃ for 5 min, and quickly chilled on ice. The dsDNA and ssDNA were used in the pull-down assay instead of RNA as described above.

### Immunofluorescence microscopy and confocal microscopy

Cells were fixed with 4% (w/v) paraformaldehyde (PFA) in PBS (pH 8.0) for 15 min and incubated with 5% (v/v) goat serum (Cedarlane, Burlington, ON, Canada), in PBS (pH 8.0) containing 0.25% (w/v) TritonX-100 for 15 min. The following primary antibodies were used: anti-LSM14A rabbit polyclonal antibody (1: 500; HPA017961, Atlas Antibodies, Stockholm, Sweden), anti-p24 mouse monoclonal antibody (1: 500; MAB7360, R&D, Minneapolis, MN, USA) and anti-ZCCHC3 mouse monoclonal antibody (1: 500; SAB1408147, Sigma-Aldrich). The following secondary antibodies were used: Alexa Fluor-647 goat anti-mouse IgG (1: 1,000; ab150115, Abcam, Cambridge, UK), Alexa Fluor-568 goat anti-rabbit IgG (1: 1,000; A-11036, Thermo Fisher Scientific) and Alexa Fluor-488 goat anti-mouse IgG (1: 1,000; A-11034, Thermo Fisher Scientific, Waltham, MA, USA). Nuclei were stained with DAPI. For the cells expressing fluorescence protein-tagged proteins, the cells were fixed with 4% (w/v) PFA in PBS (pH 8.0) and stained with DAPI. The cells were observed using a confocal laser scanning microscope (FV-3000, Olympus, Tokyo, Japan) with a 100x objective lens (NA 1.42.). The obtained images were analyzed with ImageJ (v 1.52q, NIH, Bethesda, MD, USA). A stage incubator (TOKAI HIT Corporation, Shizuoka, Japan) was used for live-cell imaging; all observations were performed at 37 °C and 5.0% CO_2_.

### Purification of virus from culture medium by centrifugation

Cells were separated from culture medium by low-speed centrifugation (700 g, 5 min, 4 ℃). The cell lysate was prepared as described in the previous section (Protein pull down assay). For concentrating the virion from the culture medium, the medium was layered onto 20% (w/v) sucrose in PBS (pH 7.4) layer and centrifuged at 20,380 g for 2 h at 4 ℃. After removing the supernatant, the pellet was resuspended with ice-cold lysis buffer (PBS containing 1% (v/v) protease inhibitor cocktail (25955-11, Nacalai Tesque) 0.25% (w/v) Triton X-100, pH 7.4). The lysates (cell and virus) were subjected to SDS-PAGE (12% (w/v) acrylamide gel) and western blot analyses using anti-EGFP antibody (1: 2,000; A-11036, Medical & Biological Laboratories Co., Ltd.), anti-p24 antibody (1:500; MAB7360, R&D), and anti-β actin antibody (1:5,000; A5441, Sigma-Aldrich).

### Flow cytometry

HeLa and TZM-bl cells were fixed with 4% (w/v) PFA in PBS (pH 8.0) for 15 min at 25 ℃. The cells were washed three times and re-suspended with PBS (pH 7.4). After filtration through nylon mesh, the cells were analyzed by a flow cytometer (LSRFortessa, BD Biosciences or Attune CytPix Flow Cytometer, Thermo Fisher Scientific). Data were analyzed using FlowJo software (vX). To confirm the expression of HA-tagged ZCCHC3 and cGAS, the cell lysate was subjected to SDS-PAGE and western blot analyses using anti-HA antibody (1:2,000, 2999S, Cell Signaling Technology).

### p24 ELISA

The culture medium of Lenti-X 293T or HEK293T cells were harvested and centrifuged at 1,500 g for 10 min to remove cell debris. The p24 amount in the medium was quantified by Lenti-X p24 Rapid Titer Kit (632200, Takara Bio Inc.) with the p24 control samples, following the manufacturer’s instructions.

### HiBiT assay

For culture supernatant of Lenti-X 293T cells transfected with the psPAX2-IN/HiBiT plasmid, the HiBiT value was measured 2 days after transfection using the Nano Glo HiBiT Lytic Detection System (N3040, Promega) as described previously^51^. The HiBiT value was converted to p24 value based on a standard curve generated with a HiBiT-containing lentiviral vector whose p24 level was already determined by p24 ELISA.

### Zcchc3 depletion

To deplete *Zcchc3*, Lenti-X 293T cells adjusted to 1.25 × 10^6^ cells per well in a 6-well plate were transfected with TriFECTa® RNAi Kit (hs.Ri.ZCCHC3.13, REF#: 107099486, IDT, Coralville, IA, USA) or non-targeting control siRNA with TransIT-X2 Dynamic Delivery System (V6100, Takara Bio Inc.) in Opti-MEM. After overnight culture, the cells were re-plated on a new 96-well plate at 2.5 × 10^4^ cells per well. The cells were cultured again overnight and subjected to the quantification of mRNA by qRT-PCR using the CellAmp Direct RNA Prep Kit for RT-PCR (Real Time) (3732, Takara Bio Inc.), One Step TB Green PrimeScript PLUS RT-PCR Kit (Perfect Real Time) (RR096A, Takara Bio Inc.), and primer pairs for *Zcchc3* (5′-CTCTCTATGCCTTCTTAAACCGA-3′ and 5′-CATCTGCACGCTACAGTTCT-3′) and *ACTB* (5′-ACAGAGCCTCGCCTTTG-3′ and 5′-C CTTGCACATGCCGGA G-3′). qRT-PCR was performed using the QuantStudio 5 Real-Time PCR System (Thermo Fisher Scientific), and the Ct values of *Zcchc3* were normalized to the mean values obtained using *ACTB* as a housekeeping gene (ΔΔCt method).

### Generation of Zcchc3 knockout cells

LentiCRISPRv2 plasmids (52961, Addgene) targeting *Zcchc3* gene were generated as follows. The following oligos (100 pmol) were mixed and heated at 95 ℃ for 5 min, followed by incubation at room temperature for 1 h for annealing, with 5′-caccgCCTGTTCCTACGCGTCTACG-3′ and 5′-aaacTAACCTCTCGGAGCCTCTGCc-3′ for sgRNA#1, and 5′-caccgCCTGTTCCTACGCGTCTACG-3′ and 5′-aaacTAACCTCTCGGAGCCTC TGCc-3′ for sgRNA#2. The mixture was 250-fold diluted with water and used for ligation with the lentiCRISPRv2 plasmid, which was predigested with Esp3I (R0734S, NEB). The solution was mixed with DNA Ligation Kit <MIGHTY Mix> (6023, Takara Bio Inc.) and used for transformation with NEB 5-alpha F′Iq Competent *E. coli* (High Efficiency) (C2992H, NEB, Ipswich, MA, USA). After the miniprep, the nucleotide sequence of the plasmid was verified by nucleotide sequencing using a primer (5′-GAGGGCCTATTTCCCATGATT-3′). The lentiCRISPRv2-Zcchc3-sgRNA#1 or lentiCRISPRv2-Zcchc3-sgRNA#2 plasmids were used for co-transfection with psPAX2-IN/HiBiT and pMD2.G plasmids on Lenti-X 293T cells using TransIT-293 Transfection Reagent. The culture supernatant was collected 2 days after transfection and used for infection on Lenti-X 293T cells. The cells were cultured for 2 weeks in the presence of 1 µg/mL puromycin (ant-pr-1, InvivoGen). The cells were single-cell-cloned using a limiting dilution method. Presence or absence of a ZCCHC3 specific band with each clone was determined by western blotting using an anti-human ZCCHC3 antibody (SAB1408147, Sigma-Aldrich).

### Reverse Transcription quantitative PCR

Lenti-X 293T cells were co-transfected with pNL4-3 with or without pCMV-HA-ZCCHC3 plasmid. At 2 days after transfection, the total RNA was extracted from cells using an RNeasy Mini Kit (74104, QIAGEN, Hilden, Germany) and QIAshredder (79656, QIAGEN,, Hilden, Germany), and subjected to RT-qPCR for quantification with the primer pairs for Gag-coding region (5′-TGTAATACCCATGTTTTCAGCA-3′ and 5′-TCTGGCCTGGTGCAATAGG-3′) and *ACTB* (5′-TCCAAATATGAGATGCGTTGTT-3′ and 5′-TGCTATCACCTCCCCTGTGT-3′). The qPCR was performed using the StepOne Plus Real-Time PCR System (Thermo Fisher Scientific). The Ct values of Gag-coding region were normalized to the mean values obtained using *ACTB* as a housekeeping gene (ΔΔCt method).

For quantifying lentiviral RNA in the virion released from the producer cells to the culture medium, RNA was purified from the medium with NucleoSpin® RNA Virus kit (U0956A, Takara Bio Inc.) by following the manufacture’s instruction and subjected to RT-qPCR with the primer pairs for EGFP (5′-CAAGCTGACCCTGAAGTTCATCTG-3′ and 5′-TTGAAGAAGTCGTGCTGCTTCATG-3′) and U5-coding region (5′-TCTGGCTAACTAGGGAACCCACTG-3′ and 5′-ACTGCTAGAGATTTTCCACACTGA C-3′). The PCR mix (15 ul) contained One step TB Green RT-PCR kit (RR096A, Takara Bio Inc.), each primer, 2 µL of nucleic acid samples and water. The primers were presented at a final concentration of 400 nM. The qPCR was performed using the StepOne Plus Real-Time PCR System (Thermo Fisher Scientific).

### Preparation of viral RNA

The plasmid carrying HIV-1 5′ LTR (454 - 634 nt, GenBank: MN989412.1), MLV 5′ LTR (1 - 207 nt, GenBank: KU324804.1), EIAV 5′ LTR (1 - 114 nt, GenBank: AF247394.1), SIV 5′ LTR (1 - 351 nt, GenBank: DQ374657.1) and Gag (935 - 1115 nt, GenBank: MN989412.1) under T7 promotor was linearized at the 3’ end of the insert, and used as a template in *in vitro* transcription reaction (MegaSscript T7 Transcription Kit (AM1333, Ambion, Waltham, MA, USA). The synthesized RNA was purified by isopropanol precipitation and quantified by measuring OD 260 nm. Short stem-loop RNA molecules derived from HIV-1 LTR (SL-1, 2, and 3) were synthesized by FasMac. The sequences were as follows: SL1, 5’-UCUCUGGUUAGACCAGAUCUGAGCCUGGGAGCUCUCUGGCUAACUAGGGA-3’; SL2, 5’-CCACUGCUUAAGCCUCAAUAAAGCUUGCCUUGAGUGCUCAAAGUAGUGU-3’; and SL3, 5’-CUAGAGAUCCCUCAGA CCCUUUUAGUCAGUGUGGAAAAUCUCUAG-3’.

### Electrophoretic mobility shift assay (EMSA)

ssRNA, dsDNA and ssDNA were prepared as described in the previous section (RNA and DNA pull down). Synthesized nucleic acids (0.1 pmol) were incubated with purified proteins (0, 0.1, 0.2, 1 pmol) in EMSA binding buffer (40 mM Tris (pH 8.0), 2 mM KCl, 1 mM MgCl_2_, 1% (w/v) NP-40, 1 mM DTT in RNase-free water) at 25 ℃ for 10 min. The samples were separated by electrophoresis using 3.5% (w/v) acrylamide/bisacrylamide gel in TBE (100 mM Tris-base, 100 mM boric acid, 2 mM EDTA), and visualized with SYBR™ Gold Nucleic Acid Gel Stain (S11494, Thermo Fisher Scientific).

### Stable cell line generation and BioID analysis

HEK293T cells were co-transfected with pDON-5neo-TurboID tagged ZCCHC3, pGP and pMD2.G at the ratio of 2:1:1 by PEI. The culture medium was collected 48 hours after transfection, centrifuged at 1,500 g for 10 min at 4 ℃ to remove cell debris, and added to the culture medium of HeLa cells. G418 was added to the medium at a final concentration of 100 μg/mL 48 hours after infection. Cells were diluted to a 96-well-plate at the concentration of 1 - 5 cells per well. Positive wells were screened using a confocal laser scanning microscope (FV-3000, Olympus) The cells were infected by lentivirus produced as described in the previous section (Plasmid transfection, virus production and collection, virus infection) for 30 min before labelling. For labelling with biotin, the cells were incubated with 500 mM biotin for 30 min. The cells were rinsed with 5 mL ice-cold HEPES-saline (20 mM HEPES-NaOH, pH 7.5, 137 mM NaCl). Another 2 mL HEPES-saline was added to the dish to collect the cells with scraping. The cells were harvested by centrifugation at 800 g for 3 min at 4 ℃ and re-suspended with 500 µL Guadinine-TCEP buffer (8 M Guadinine, 100 mM HEPES-NaOH, pH 7.5, 10 mM tris(2-carboxyethyl)phosphine hydrochloride, 40 mM chloroacetamide). Proteins were extracted and digested with trypsin followed by enrichment of biotinylated peptides as described previously^67^. LC-MS/MS analysis of the biotinylated peptides was performed on an EASY-nLC 1200 UHPLC connected to an Orbitrap Fusion mass spectrometer (Thermo Fisher Scientific) as described previously^67^. Raw data were directly analyzed against the SwissProt database restricted to *Homo sapiens* using Proteome Discoverer version 2.5 (Thermo Fisher Scientific) with Sequest HT search engine. The search parameters were as follows: (a) trypsin as an enzyme with up to two missed cleavages; (b) precursor mass tolerance of 10 ppm; (c) fragment mass tolerance of 0.6 Da; (d) carbamidomethylation of cysteine as a fixed modification; and (e) acetylation of protein N-terminus, oxidation of methionine, and biotinylation of lysine as variable modifications.

### Luciferase assay

Cells infected with a luciferase-encoding virus were lysed 2 days after infection with a Bright-Glo Luciferase Assay System (E2620, Promega) and the luminescent signal was measured using a GloMax Explorer Multimode Microplate Reader (Promega).

### Statistics

The total number of independent experiments in each analysis is described in the figure legends. All statistical analyses were evaluated by an unpaired, two-tailed Student’s *t*-test unless otherwise indicated. *p* ≤ 0.05 were considered statistically significant. The tests were performed using Prism 9 software v9.1.1 (GraphPad Software Inc., Boston, MA, USA).

### Protein structure predictions

The three-dimensional protein structure of human ZCCHC3 (NM_033089.7) was predicted using ColabFold (v.1.5.2: AlphaFold2 using MMseqs2) (https://colab.research.google.com/github/sokrypton/ColabFold/blob/main/AlphaFold2.ipynb).

### Protein-RNA binding predictions

The protein-RNA binding was predicted busing catRAPID omics (v2.0) (http://s.tartaglialab.com/page/catrapid_omics2_group). The amino acids sequence of ZCCHC3, the nucleotide sequences of HIV-1 LTR RU5 region and MLV LTR used for the prediction could be found in NCBI (ZCCHC3: NM_033089.7; HIV-1 LTR RU5 region: MN989412.1; and MLV LTR: KU324804.1).

## Data availability statement

This study includes no original code deposited in external repositories. Source data are provided with this paper. Any additional information required to reanalyze the data reported in this study is available from the corresponding author upon request.

## Acknowledgements

The following reagents were obtained through the NIH HIV Reagent Program, Division of AIDS, NIAID, NIH: Anti-Human Immunodeficiency Virus 1 (HIV-1) p24 Monoclonal (183-H12-5C), ARP-3537, contributed by Dr. Bruce Chesebro and Kathy Wehrly. This research was supported by Japan Agency for Medical Research and Development (AMED) under grants JP22fk0410033 (to A.S.), JP21fk0108465 (to A.S.), JP22jk0210039 (to A.S.), JP22wm0325009 (to A.S. and S.H.Y.) and JP22fk0410047 (to A.S. and S.H.Y.); JSPS Grants-in-Aid for Scientific Research on Innovative Areas (19H04830 to S.H.Y.), JSPS Grant-in-Aid for Scientific Research (A) (23H00369 to S.H.Y.), JSPS Grant-in-Aid for Scientific Research (C) (19K06382 to A.S.), and JSPS Grant-in-Aid for Scientific Research (B) (22H02500 to A.S.); The Ito Foundation Research Grant R4 (to A.S.); Grant for Joint Research Projects of the Research Institute for Microbial Diseases, Osaka University (to A.S.); and the Joint Usage and Joint Research Programs, Institute of Advanced Medical Sciences, Tokushima University (to H.K. and S.H.Y.). We thank S. Dodo, Y. Shibatani, T. Nishiuchi, M. Kumeta, Y. Yu, and A. Jimpo for technical assistance.

## Author contributions

Experiments were designed by B.Y., A.S., J.L., P.S., and S.H.Y. Plasmids construction, recombinant protein purification, protein pull-down assay, RNA pull-down assay, EMSA, RNA preparation, reverse transcription quantitative PCR, and stable cell generation were performed by B.Y. and P.S. Cell culture and transfection, confocal fluorescence microscopy, flow cytometry was performed by B.Y., Y.L.T., E.P.B., S.H.Y., and A.S. BioID assay was performed by B.Y. and H.K. ZCCHC3 knockout cells generation and HiBiT assay were performed by Y.L.T. and A.S. Data analysis was completed by B.Y., S.H.Y., and A.S. Manuscript writing, figure design, and editing was done by B.Y., H.K. S.H.Y and A.S. S.H.Y and A.S. supervised and funded the project.

## Additional information

Supplementary Information is available for this paper.

Correspondence and requests for materials should be addressed to Akatsuki Saito and Shige H. Yoshimura.

Reprints and permissions information is available at www.nature.com/reprints.

---

**Extended Data Fig. 1.**
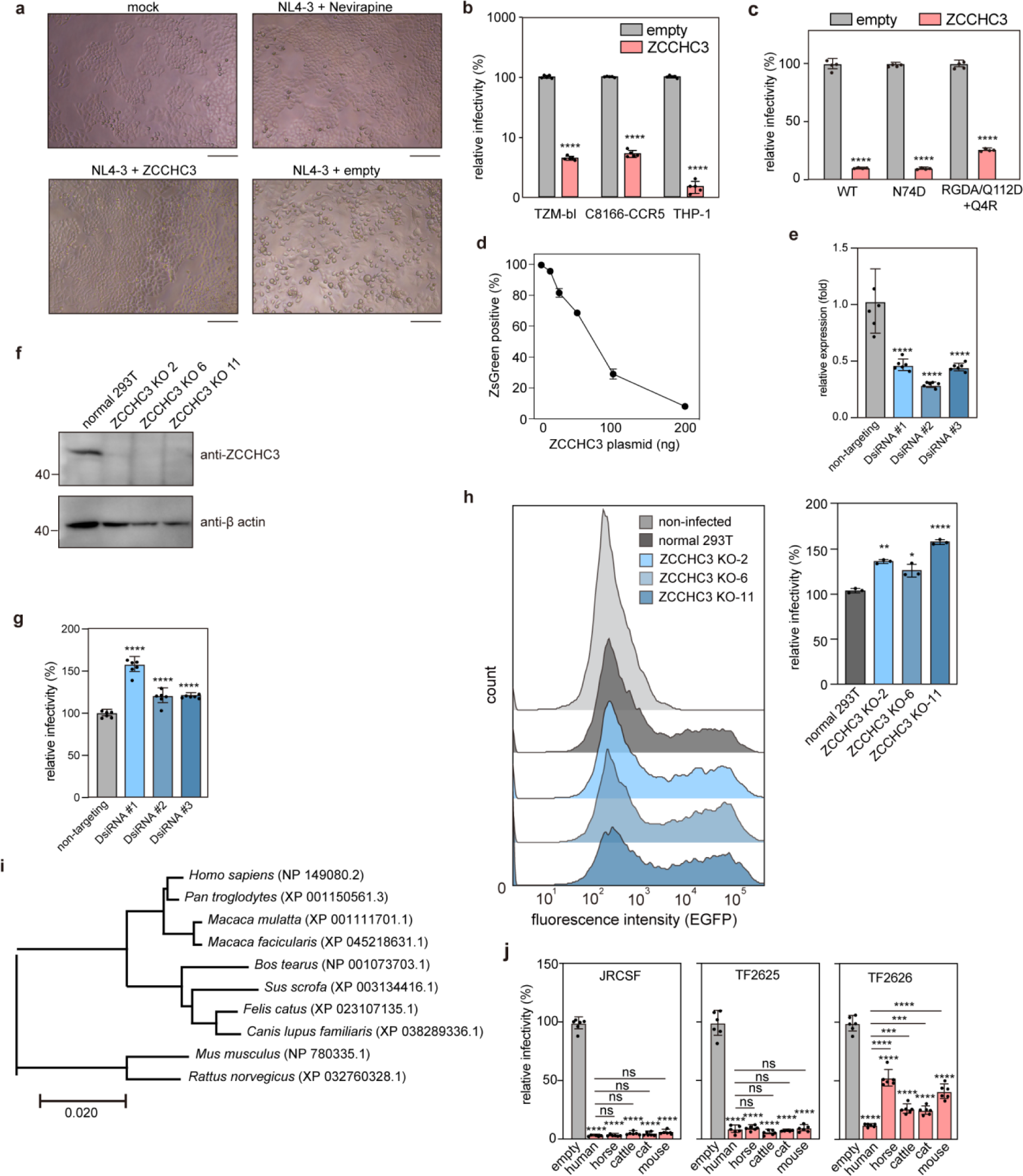
Additional data on the effect of ZCCHC3 on viral infection. **a**, TZM-bl cells were infected with NL4-3 viruses produced in the presence or absence of a HA-ZCCHC3 expression plasmid. The cells were observed by an optical microscopy 2 days after infection. TZM-bl cells infected with NL4-3 in the presence of Nevirapine served as a control (n =1). Scale bar, 50 µm. **b**, Antiviral effect of ZCCHC3 is Env-independent. pMSMnG plasmid was introduced into Lenti-X 293T cells with or without a ZCCHC3 expression plasmid. VSV-G-pseudotyped viruses were collected 2 days after transfection and used to infect the indicated cells. Infectivity on TZM-bl cells was determined as in (Fig. 1c); otherwise, it was determined by flow cytometry as a percentage of GFP-positive cells 2 days after infection. Values relative to those for cells harbouring empty vector are shown as the mean ± standard deviation (n = 5). **c**, Sensitivity of CA mutants to ZCCHC3. pMSMnG plasmid encoding the indicated HIV-1 CA variant was introduced into Lenti-X 293T cells with or without a ZCCHC3 expression plasmid. Culture supernatant was collected 2 days after transfection and used to infect C8166-CCR5 cells. Infectivity was determined by flow cytometry as in (Extended Data Fig. 1b). Values relative to those for cells harbouring empty vector are shown as the mean ± standard deviation (n = 6). **d**, Dose-dependent suppression of lentiviral infectivity by ZCCHC3. Lenti-X 293T cells were co-transfected with EGFP-expressing lentiviral vectors together with different amounts of HA-ZCCHC3 expression vector. Culture supernatant was collected 2 days after transfection and used to infect MT-4 cells. Cell fluorescence was measured by flow cytometry 2 days after infection. The mean and standard deviation values from quadruplicate measurements are shown. **e**, Knock down of Z*cchc3*. *Zcchc3* mRNA in Lenti-X 293T cells transfected with DsiRNA targeting *Zcchc3* or control DsiRNA was quantified by RT-qPCR. The mean and standard deviation values are shown (n = 6). **f**, Western blot analysis of ZCCHC3-KO cells. Three KO cell lines were analyzed by western blotting using an anti-ZCCHC3 antibody, with β-actin as a loading control. A representative image from three independent experiments is shown. **g**, Effect of *Zcchc3* knockdown on viral infectivity on target cells. Lenti-X 293T cells transfected with DsiRNA targeting *Zcchc3* or control DsiRNA were infected with a lentiviral vector encoding the luciferase reporter protein. Infectivity was quantified as relative light units of luciferase and is presented as the mean and standard deviation (n = 6). **h**, Infectivity of lentivirus produced from normal and ZCCHC3-KO cells, analyzed by flow cytometry (left). Positive cells were counted (right). The mean and standard deviation values are shown from three independent experiments. **i**, Phylogenetic tree of mammalian ZCCHC3 proteins. The tree was constructed using the maximum-likelihood method. **j**, Anti-HIV-1 effect of mammalian ZCCHC3. A plasmid encoding the indicated virus was introduced into Lenti-X 293T cells with or without mammalian HA-ZCCHC3 expression plasmid. Culture supernatant was prepared, and TZM-bl cells infected and analyzed as in (Fig. 1c). Values relative to that of human ZCCHC3 are shown as the mean and standard deviation (n = 6). In **b,c,h**, differences were examined by a two-tailed, unpaired Student’s *t*-test; \*\*\*\**p* < 0.0001, \*\**p* < 0.01, \**p* < 0.05. In **e,g** and **j** differences were examined by one-way ANOVA, followed by Tukey’s test; \*\*\*\**p* < 0.0001, \*\*\**p* < 0.001, ns *p* < 0.01.

**Extended Data Fig. 2.**
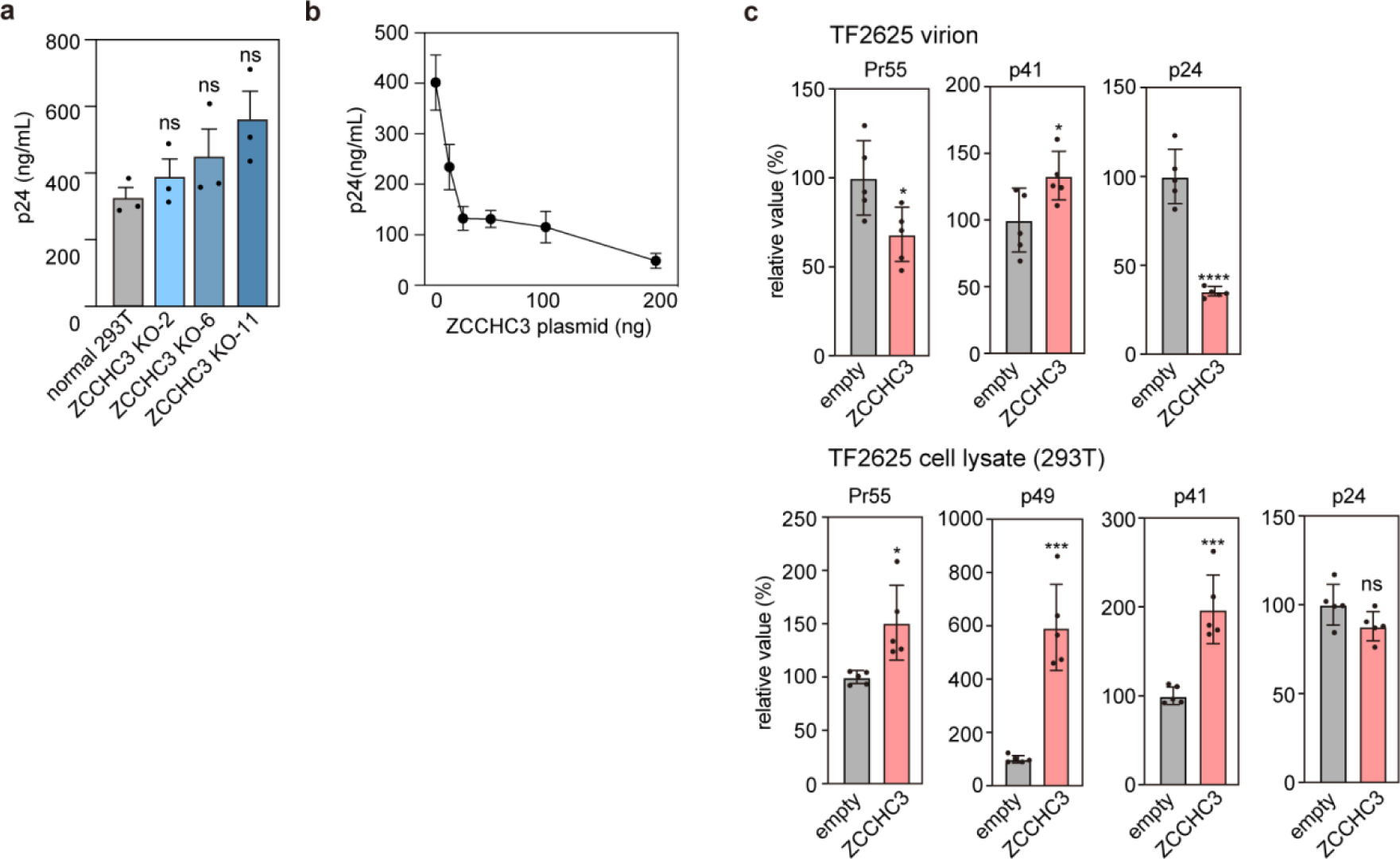
Additional data on the effect of ZCCHC3 on viral production. **a,b**, The amount of HIV-1 released into the culture medium described in Extended Data Fig. 1h (**a**) and Extended Data Fig. 1c (**b**), quantified by p24 ELISA. The mean and standard deviation values from three independent experiments are shown. **c**, Quantified band intensity from blots shown in Fig. 2b. Relative intensity of pr55, p49, p41, and p24 bands in the virion lysate (upper), or pr55, p49, p41, and p24 bands in the cell lysate (lower) of pTF2625-transfected cells was compared in the presence or absence of ZCCHC3 during virion production (n = 5). In **a,c**, differences were examined by a two-tailed, unpaired Student’s *t*-test. \*\*\*\**p* < 0.0001, \*\*\**p* < 0.001, \**p* < 0.05; ns, *p* ≥ 0.05.

**Extended Data Fig. 3.**
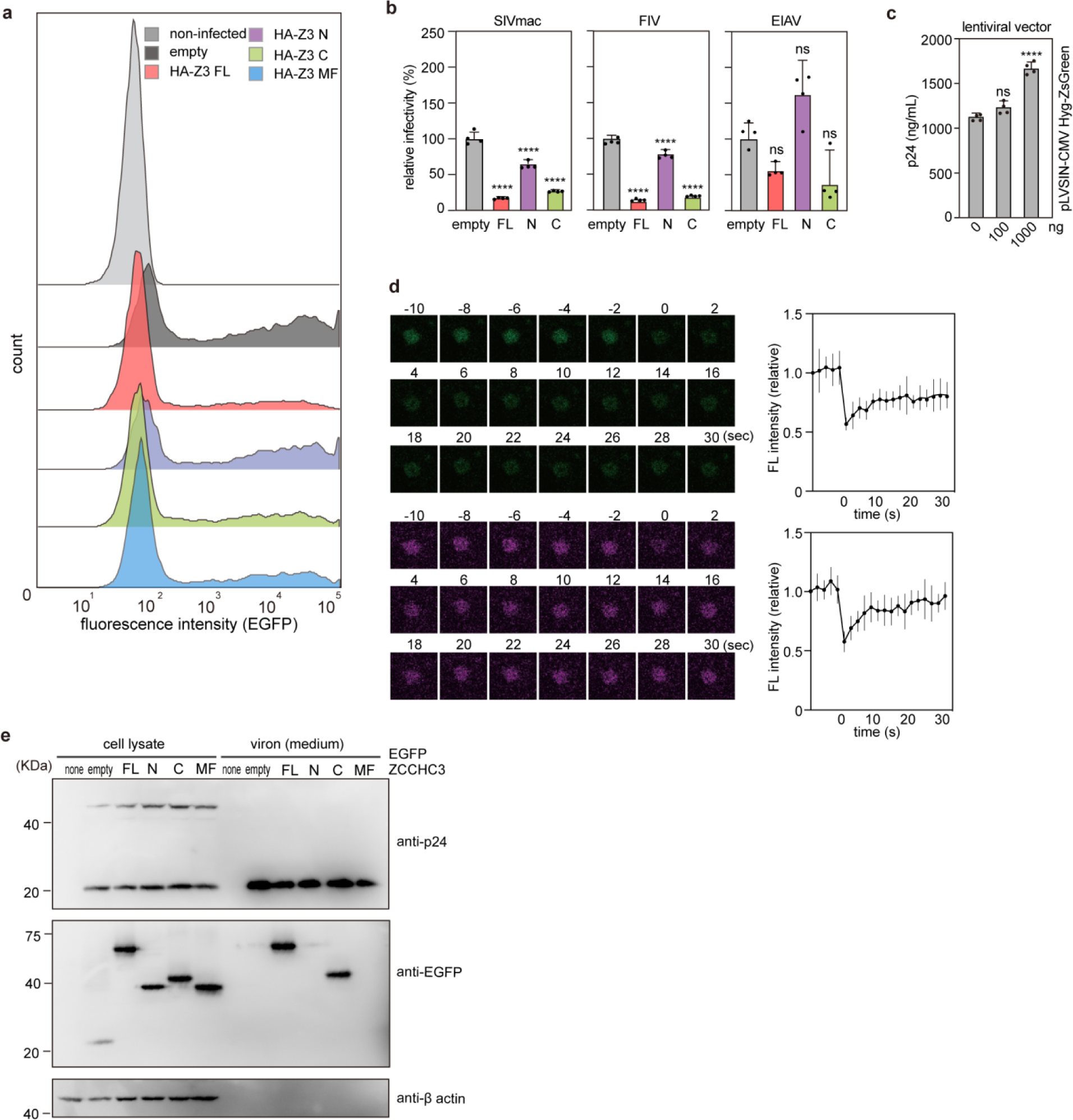
Additional data on ZCCHC3 binding to GagNC via C-terminal ZnFs. **a**, Changes of lentiviral infectivity upon ZCCHC3 loading. HeLa cells were infected with a p24-normalized amount of lentiviruses carrying a ZCCHC3 fragment (FL, N, C, or MF). Infectivity was analyzed based on the expression of a viral gene (EGFP) using flow cytometry. **b**, ZCCHC3 C fragment suppresses retroviral infection. Lenti-X 293T cells were co-transfected with plasmids for generating SIVmac, FIV, or EIAV lentiviral vectors in the presence of a ZCCHC3 fragment (FL, N, or C), and the resultant viruses were used to infect MT-4 cells. Infectivity was determined as relative light units 2 days after infection. Values relative to those of cells without ZCCHC3 expression are shown as the mean ± standard deviation (n = 5). **c**, Lenti-X 293T cells were co-transfected with 1000 ng of psPAX2-IN/HiBiT plasmid and different amounts of pLVSIN-CMV Hyg-ZsGreen vector. p24 concentration (converted from HiBiT value) in the culture supernatant was determined 2 days after transfection. Values are shown as the mean ± standard deviation (n = 4). **d**, FRAP analysis of a condensate formed by HIV-1 Gag NCp7 in HeLa cells. The foci were bleached, and the fluorescence intensity in the area was measured. Data are shown as the mean (bold solid line) and standard deviation (vertical lines) (n = 10). **e**, Incorporation of different ZCCHC3 fragments into the lentiviral virion. A plasmid encoding EGFP-tagged ZCCHC3 FL, N, C, or MF was introduced into HEK293T cells together with lentiviral plasmids (pLV-SIN-Hyg, psPAX2, pMD2.G). Virions released into the culture medium were harvested by centrifugation and analyzed by immunoblotting with anti-GFP, anti-p24, and anti-β-actin antibodies. In **b,c**, differences were examined by one-way ANOVA, followed by Tukey’s test. \*\*\*\**p* < 0.0001; ns, *p* ≥ 0.05.

**Extended Data Fig. 4.**
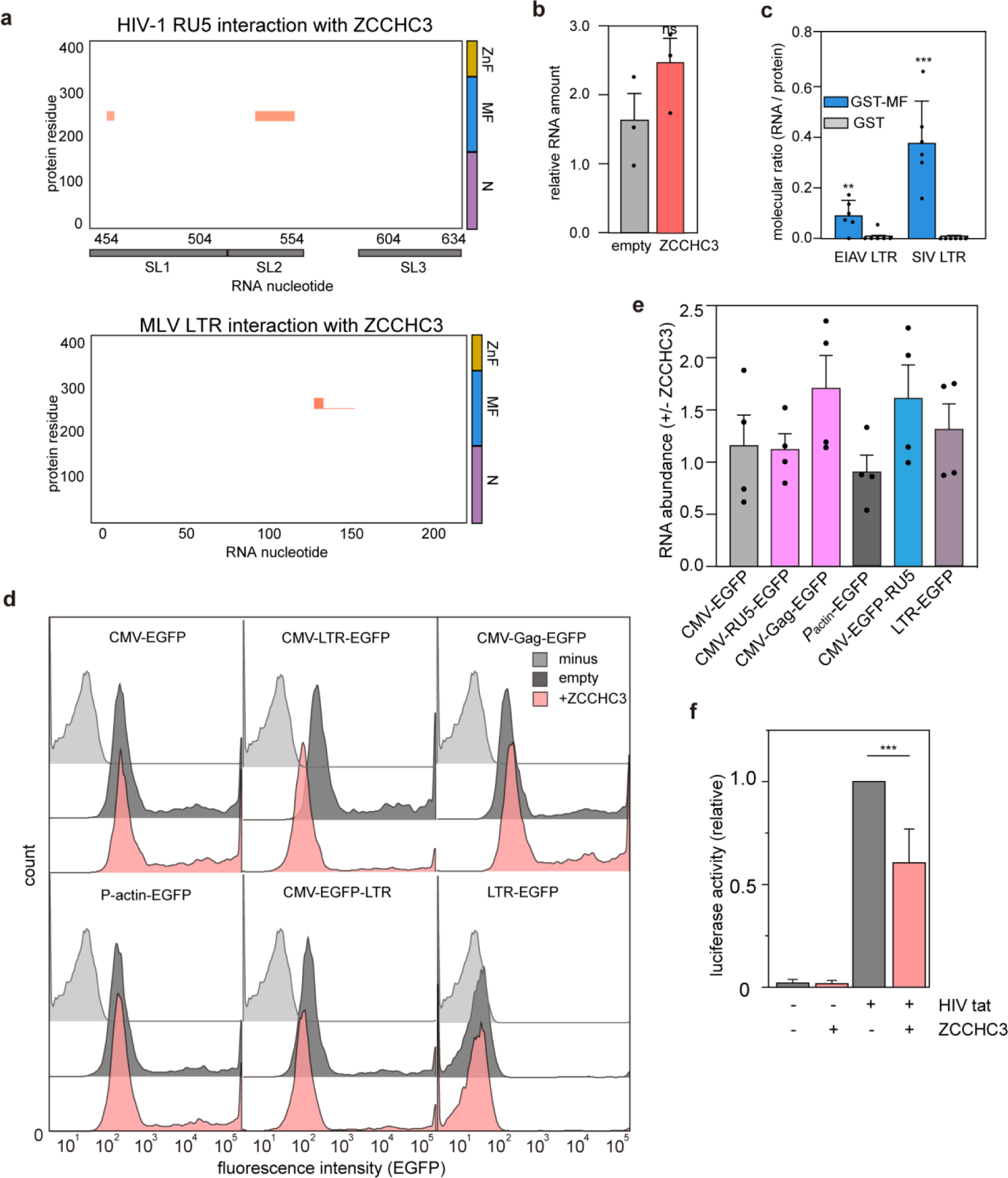
Additional data on ZCCHC3 binding to retroviral RNA. **a**, Binding propensity of ZCCHC3 to HIV1-1 LTR (R-U5) RNA sequence and MLV LTR sequence, predicted by catRAPID. The nucleotide numbers of HIV-1 (GenBank: MN989412.1) and MLV genomes (GenBank: KU324804.1) are plotted on the x-axis. The HIV-1 SL1, SL2, and SL3 positions are depicted along the x-axis. ZCCHC3 N, MF, and ZnF positions are depicted along the y-axis. **b**, Effect of ZCCHC3 on the expression of viral genes. pNL4-3 plasmid was introduced into Lenti-X 293T cells together with a vector encoding HA-ZCCHC3. Total RNA was purified, and viral RNA was quantified by RT-qPCR and normalized to β-actin mRNA expression level. The mean and standard deviation from three independent experiments are shown. **c**, ZCCHC3 MF domain binding to EIAV and SIV LTR RNA. RNA pull-down assay was performed using the ZCCHC3 MF fragment and EIAV LTR and SIV LTR, as in Fig. 4b. The mean and standard deviation values from three independent experiments. **d**. Representative traces of the reporter genes expression level (Fig. 5g). **f**, Effect of ZCCHC3 on viral gene expression in a proviral state. TZM-bl cells carrying a luciferase gene flanked by 5′- and 3′-LTRs were transfected with a plasmid encoding HIV-1 Tat with or without HA-ZCCHC3. Luciferase activity in the cell lysate was then measured. The data are presented as the mean ± standard deviation from four independent experiments. In **b–d** and **f,** differences were examined by a two-tailed, unpaired Student’s *t*-test. \*\*\**p* < 0.001, \*\**p* < 0.01, \**p* < 0.05; ns, *p* ≥ 0.05.

**Extended Data Fig. 5.**
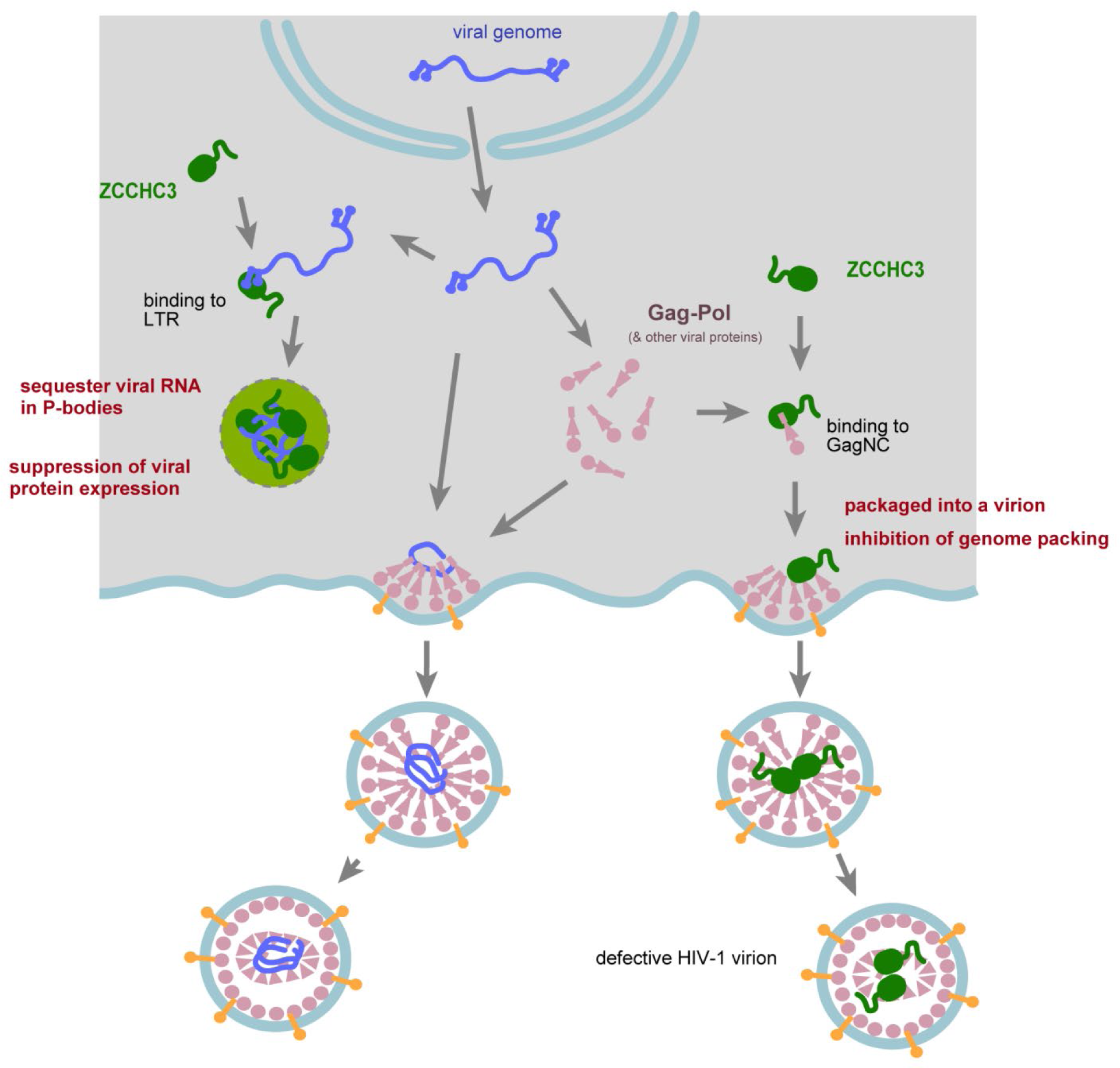
Proposed mechanism of HIV-1 suppression by ZCCHC3. The 5′ LTR region of nuclear-exported HIV-1 genomic RNA is recognised by the ZCCHC3 MF, and sequestered in the P-bodies, which impairs virion maturation. ZCCHC3 also binds to HIV-1 Gag NCp7, which leads to ZCCHC3 incorporation into the virion and promotes the antiviral function of ZCCHC3 during subsequent infection.

